# Ex vivo maturation of the malaria parasite egress protease SERA6 aids pathway dissection and inhibitor development

**DOI:** 10.64898/2026.01.29.702321

**Authors:** Chrislaine Withers-Martinez, Zahie Taha, Christine R Collins, Fiona Hackett, Michele SY Tan, Christelle Soudy, Dhira Joshi, Joanna Redmond, Benedict Davies, Sarah Maslen, Mark Skehel, Roger George, Svend Kjaer, Michael J Blackman

## Abstract

Release (egress) of malaria parasites from host red blood cells (RBC) is a protease-dependent process involving breakdown of the RBC cytoskeleton by a parasite cysteine protease-like protein called SERA6. In the penultimate step of the egress cascade, SERA6 undergoes autoproteolytic maturation triggered upon cleavage by a serine protease called SUB1 and requiring interactions between SERA6 and fragments of another parasite protein called MSA180. Egress can be blocked by treatment of intraerythrocytic parasites with small molecules that prevent the autocatalytic SERA6 maturation step, suggesting that SERA6 is a druggable target. Here we describe the development of a cell-free in vitro system that recapitulates SERA6 maturation. We use the assay to confirm the strict requirement for MSA180 in SERA6 maturation by SUB1 and to show that these 3 components are sufficient for SERA6 maturation. Using a synthetic peptide substrate based on a predicted autocatalytic cleavage site we demonstrate that the fully mature SERA6 is an active proteolytic enzyme and we validate improved small molecule inhibitors of SERA6. Our lead inhibitory compound efficiently blocks egress of asexual blood stage parasites, confirming SERA6 as a new potential antimalarial drug target.

## Introduction

Malaria continues to exact extensive morbidity and mortality across the globe, with children below the age of 5 years and pregnant women at particular risk of severe disease (World Health Organization World Malaria Report 2025). Despite recent promising progress in the development of protective vaccines [1], chemotherapy remains crucial for malaria control. There is therefore widespread concern over the continuing spread of resistance of the causative agents, protozoan parasites of the genus *Plasmodium*, to mainstay artemisinin-based antimalarial drugs [2, 3]. An improved understanding of fundamental parasite biology, especially that of the species causing the majority of severe disease, *Plasmodium falciparum*, is essential for the development of new target-based drugs, as well as for elucidating the mode of action of novel drug-like antimalarial compounds discovered through phenotypic screens [4, 5].

Clinical malaria results from asexual replication of the parasite in circulating red blood cells (RBC). Parasite growth within the RBC, enabled through catabolism of RBC haemoglobin in a specialised parasite digestive vacuole, culminates in the formation of a mature multinucleated form called a schizont. Multiple daughter merozoites bud from the intracellular schizont, then rupture of the infected cell allows egress of the merozoites which rapidly invade fresh RBC to initiate a new replicative cycle. Egress involves several characteristic morphological and structural transitions, including first permeabilization (‘poration’) of the intraerythrocytic parasitophorous vacuole (PV) within which the schizont resides [6], then rounding up of the PV followed by its rupture with a concomitant microscopically discernible increase in mobility of the intraerythrocytic merozoites. This is rapidly followed by poration of the RBC membrane, then final breakage of the RBC membrane to allow merozoite dissemination in an explosive manner [7–13]. Several lines of genetic and biochemical evidence have shown that egress is controlled through a proteolytic cascade triggered by the activation of a cGMP-regulated parasite kinase called PKG [14]. PKG activity is itself dependent upon a guanylyl cyclase (GCα) [15] as well as a serine-threonine phosphatase called PP1 [16], whilst PKG appears to function in coordination with a multi-pass membrane protein called ICM1 and a calcium-dependent protein kinase, CDPK5 [17–19] PKG activation leads to a parasite subtilisin-like serine protease called SUB1 being discharged from specialised merozoite subcellular organelles called exonemes into the lumen of the PV [20, 21]. There, SUB1 encounters a cysteine protease-like parasite molecule called SERA6 [22]. Precise SUB1-mediated cleavage of the SERA6 precursor at two defined sites releases a central SERA6 module called p65, comprising an N-terminal prodomain fused to the papain-like putative catalytic domain. A rapid autocatalytic cleavage event then releases the SERA6 prodomain (referred to as p40) from p65, generating the terminal papain-like SERA6 domain, SERA6 PAP. This interacts with at least 2 fragments of a second PV-resident parasite protein called malarial SERA activator 180 (MSA180) which is also cleaved by SUB1 [23]. Formation of the SERA6 PAP-MSA180 complex results in rapid degradation of the RBC cytoskeletal component β-spectrin, in turn leading to destabilisation and rupture of the RBC membrane to allow merozoite egress [13, 23]. Egress also involves at least one PV-resident phospholipase activity called LCAT [24], although the precise role of this enzyme in the pathway is unclear. Genetic ablation and/or pharmacological inhibition of any of these egress mediators, or of an aspartic protease called plasmepsin X (PMX) required for SUB1 maturation [25–28], modulates, delays or prevents egress (Fig 1). The key role of SERA6 maturation in the final stages of egress is also strongly supported by evidence that small molecules that inhibit the autocatalytic p65-to-PAP maturation step selectively block β-spectrin cleavage as well as RBC membrane rupture and egress, but do not prevent rounding up or rupture of the PV, nor the temporally associated poration of the RBC membrane (all of which are SERA6 independent) [29, 30] [23]. These observations suggest that SERA6 could be a target for a new generation of protease-inhibitor based antimalarial drugs.

**Figure 1.**
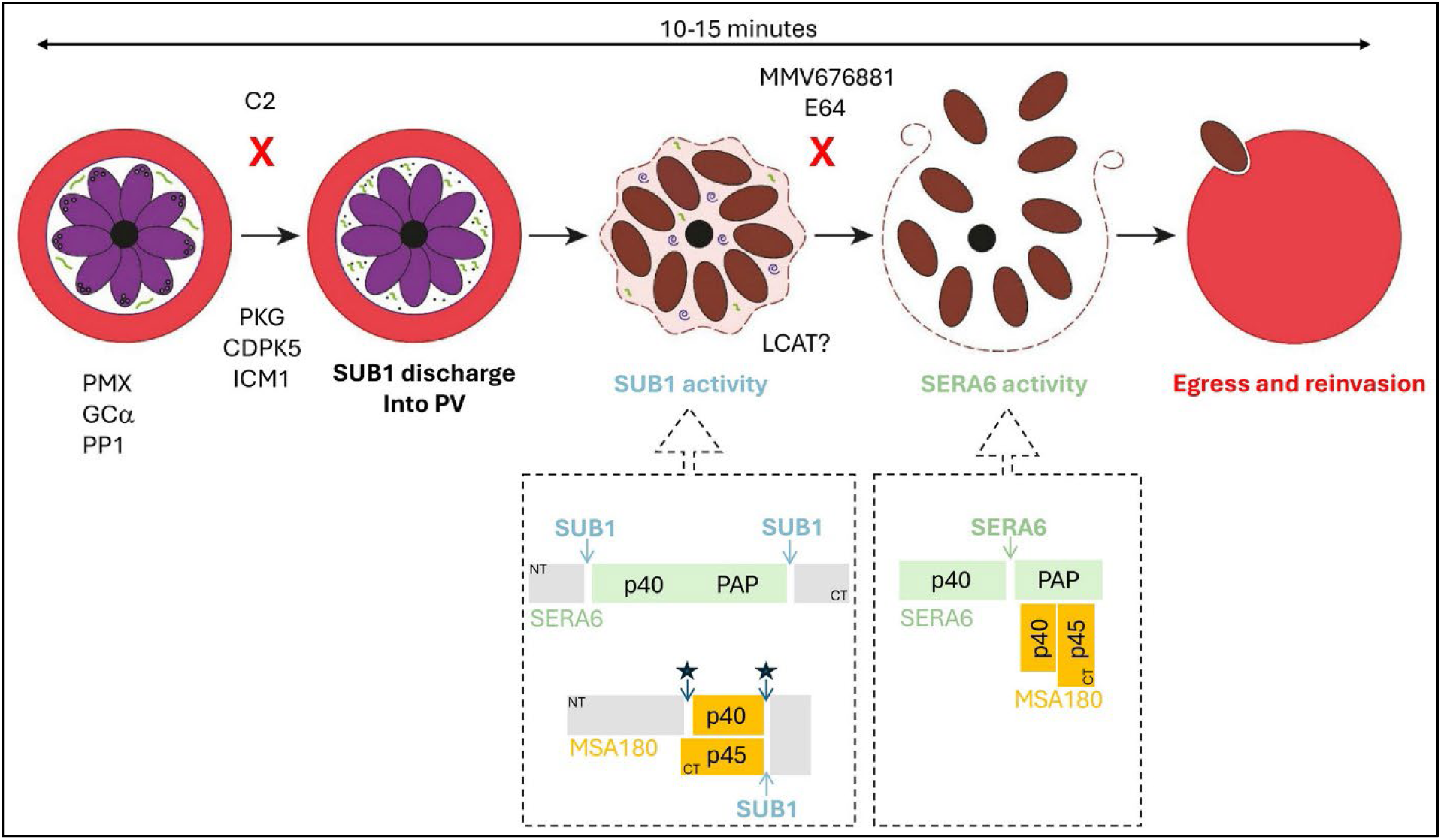
Model of the enzyme pathway that regulates malaria parasite egress from the host RBC. Key molecular players and their relative position in the egress pathway are shown. PMX is required for maturation of SUB1, which is stored in specialized secretory organelles called exonemes. Activity of the cGMP-dependent kinase PKG is dependent upon GCα and PP1, and PKG functions in coordination with CPDK5 and ICM1 to regulate discharge of active SUB1 into the PV. Processing activities of SUB1 and SERA6 are detailed in the two dashed boxes. SUB1 cleaves SERA6 at two sites to release the p65 intermediate composed of the prodomain p40 and its cognate catalytic papain-like domain PAP. SUB1 also processes MSA180, releasing a p45 C-terminal fragment and a larger p40 domain. These events are followed by autocatalytic maturation of SERA6 to the terminal PAP domain and additional processing of MSA180 (starred). SERA6 PAP and MSA180 p40 and p45 domains interact to form a mature complex that is involved in cleavage of β-spectrin, leading to disruption of the RBC membrane to enable merozoite egress. Compound 2 (C2) prevents egress by inhibiting PKG activity, whilst the nitrile MMV676881 and the epoxide E64 block the autocatalytic step of SERA6 maturation. The phospholipase LCAT is also required for normal egress. NT, N-terminus; CT, C-terminus.

Despite this detailed picture of SERA6 maturation and its role in egress, it is unclear whether parasite components additional to SUB1 and MSA180 are required for SERA6 maturation. Furthermore, a direct demonstration that mature SERA6 PAP is an active protease capable of cleaving peptide-based substrates in an enzymatic manner *in trans* is lacking. Here, using parasite-derived SERA6 and MSA180 together with recombinant SUB1 we describe the development of a cell-free in vitro system that recapitulates SUB1-dependent SERA6 maturation. We use the assay to: (1) define the minimal requirements for SERA6 maturation; (2) to show for the first time that the fully processed SERA6 PAP is an enzymatically active protease; and (3) to validate improved small molecule inhibitors of SERA6, confirming SERA6 as a new potential antimalarial drug target.

## Results

### *Ex vivo* maturation of SERA6 in cell-free parasite extracts using recombinant SUB1

Maturation of SERA6 in the minutes leading up to egress is initiated by SUB1-catalysed cleavage of the SERA6 precursor within the PV to release the central p65 module, which comprises an N-terminal prodomain fused to the papain-like putative catalytic domain [23, 31]. An autocatalytic cleavage event then releases the SERA6 prodomain p40, generating the free terminal SERA6 PAP, thought to be the active form of SERA6.

Formation of SERA6 PAP and its recruitment of fragments of MSA180 is thought to be important for the β-spectrin cleavage that facilitates lysis of the host RBC cytoskeleton and membrane to allow merozoite egress. Recombinant, enzymatically-active SUB1 has been available for some time [32, 33]. In contrast, extensive previous attempts by us and others to generate correctly-folded recombinant full-length (FL) SERA6, FL MSA180 or SERA6 PAP have failed, hindering efforts to further define the mechanism of and requirements for SERA6 maturation. To overcome this hurdle, we set out to develop an assay that exploits native SERA6 and MSA180 isolated from parasites cultured *in vitro* in human RBC.

The non-ionic glycoside detergent saponin readily permeabilizes RBC membranes and the PV membrane, but does not rupture the blood stage malaria parasite plasma membrane [34]. On this basis, saponin has been widely used for the preparation of parasite extracts enriched in soluble PV proteins, including FL SERA6 (e.g. [22]). We reasoned that a saponin extract of mature intraerythrocytic schizonts arrested prior to egress with the PKG inhibitor 4-[7-[(dimethylamino)methyl]-2-(4-fluorphenyl)imidazo[1,2-α]pyridine-3-yl]pyrimidin-2 amine (compound 2, C2) should contain FL SERA6 as well as all the components required for SERA6 maturation except for SUB1, which remains sequestered in exonemes in C2-arrested parasites [21]. We predicted that addition of exogenous recombinant SUB1 (rSUB1) to such *ex vivo* extracts should trigger maturation of the endogenous FL SERA6. To test this notion, we generated extracts of mature, C2-arrested schizonts using a saponin-containing buffer. To enable downstream analysis, for this work we used the previously-described transgenic *SERA6-mTAP:loxP P. falciparum* line [23], which expresses fully functional SERA6 possessing an internal haemagglutinin (HA) epitope tag. The schizont extracts were then supplemented *in vitro* with rSUB1 and the fate of the endogenous FL SERA6 monitored. As shown Fig 2A, this resulted in the expected time-dependent conversion of FL SERA6 to its mature PAP form, via the p65 intermediate. Importantly, conversion of p65 to PAP in the rSUB1-treated saponin extracts was blocked by the presence of the epoxide E64 (Fig 2B), a potent inhibitor of the autocatalytic step of SERA6 maturation and egress when used to treat intraerythrocytic parasites [13, 23]. These results indicated that it is possible to reconstitute the entire SUB1-dependent SERA6 maturation pathway in a cell-free *in vitro* system and that the autocatalytic p65-to-PAP step of SERA6 maturation under such conditions remains susceptible to small molecule inhibitors.

**Figure 2.**
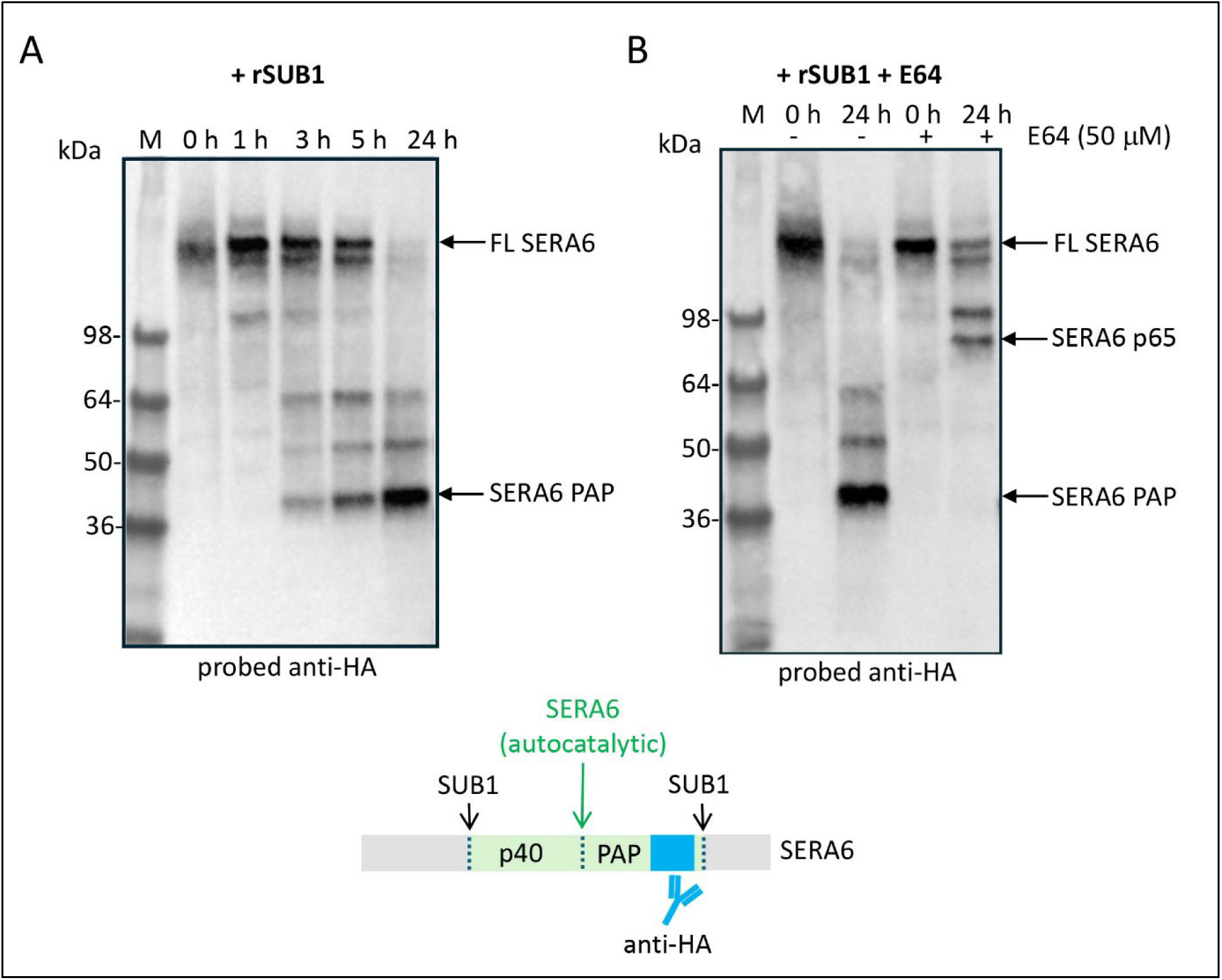
*Ex vivo* maturation of parasite-derived SERA6 using recombinant SUB1. (A) Time-course analysed by western blot showing conversion of full length SERA6 (FL SERA6) in schizont saponin extracts to the terminal PAP form by recombinant SUB1 (rSUB1), added at time 0 h. Conversion to SERA6 PAP was complete by 24 h. (B) A similar time course showing that supplementing the saponin extracts with the epoxide cysteine protease inhibitor E64 prevents the autocatalytic p65-to-PAP maturation step. Positions of molecular mass marker proteins (M) are indicated. Bottom, schematic (not to scale) showing the relative positions of the known SUB1 processing sites within FL SERA6 as well the putative autocatalytic processing site that converts p65 to p40 and PAP. Also indicated is the position of the internal HA epitope tag expressed by the transgenic *SERA6-mTAP:loxP* parasites used as a source of the saponin extracts used in these experiments.

### SUB1 and MSA180 are both necessary and sufficient for complete ex vivo maturation of SERA6

The assay described above exploited as a source of FL SERA6 unfractionated saponin extracts, which contain a broad repertoire of PV proteins. Whilst in previous work we have shown through gene disruption experiments that both SUB1 and MSA180 are required for SERA6 maturation [23, 13], it has not been possible to address whether these are sufficient or whether additional components are required. We have previously described generation of the transgenic *MSA180-HA3:loxP P. falciparum* line [23] which expresses an epitope-tagged form of MSA180. We reasoned that we could take advantage of this line as well as the *SERA6-mTAP:loxP* parasite line to independently isolate the respective full-length MSA180 and SERA6 proteins from saponin extracts using affinity-purification, then recombine the isolated proteins in the presence of rSUB1 to test whether these three components alone are sufficient for SERA6 maturation. As shown in Fig 3A and Fig 3B, the endogenous tagged SERA6 and MSA180 proteins could indeed be successfully isolated by adsorption onto anti-HA antibody beads followed by gentle elution from the beads using HA peptide. In the absence of added MSA180, purified FL SERA6 was converted by rSUB1 only to the p65 intermediate form, with no detectable formation of PAP (Fig 3C). In contrast, supplementing the eluted, reconstituted proteins with rSUB1 resulted in complete conversion of FL SERA6 to PAP (Fig 3D). FL MSA180 was also converted by rSUB1 in these experiments to a smaller product p45, consistent with the presence of at least one SUB1 cleavage site within MSA180 as previously shown [23]. Importantly, in the absence of added rSUB1 both FL SERA6 and FL MSA180 were predominantly stable over the period of incubation, with no detectable conversion to smaller products. The observed rSUB1-dependent maturation of SERA6 and MSA180 in the *ex vivo* system therefore mimicked the *in vivo* process. Taken together with our previous genetic data, these results show that SUB1 and MSA180 are both necessary and sufficient for complete SERA6 maturation to PAP, with no requirement for any other parasite component(s).

**Figure 3.**
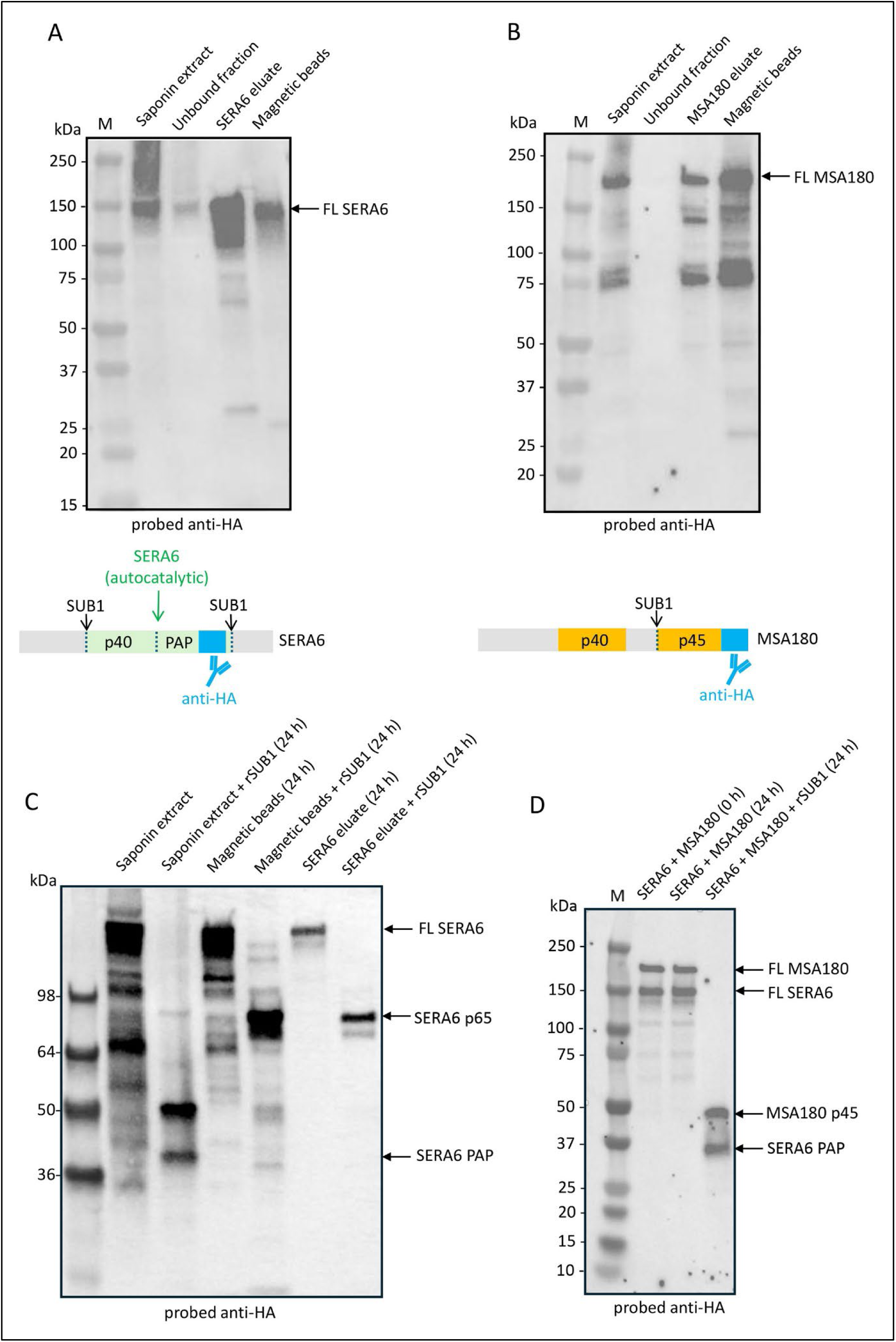
Maturation of SERA6 requires only SUB1 and MSA180. (A) and (B). Western blot analysis of the affinity purification of HA-tagged FL SERA6 (A) and FL MSA180 (B) using anti-HA antibody magnetic beads. Bound proteins were eluted with HA-peptide. Lanes labelled ‘Magnetic beads’ contain proteins remaining immobilised after elution with HA-peptide, then released from the beads by heating in SDS sample buffer. (C) Western blot analysis showing that SERA6 in the saponin extract is fully converted to the PAP form in the presence or rSUB1 and that addition of rSUB1 to affinity-purified HA-tagged FL SERA6 alone, either still immobilised on magnetic beads or following elution, resulted only in conversion to the p65 form. (D) Western blot showing the results of incubation of affinity-purified FL SERA6 plus FL MSA180 with or without added rSUB1. The purified proteins remained intact throughout the 24 h incubation in the absence of rSUB1, whereas incubation with rSUB1 resulted in their complete conversion to SERA6 PAP and the P45 fragment of MSA180 respectively. Positions of migration of molecular mass marker proteins are indicated (M). Middle, schematic (not to scale) showing the positions of SUB1 processing sites and the HA epitope tags within SERA6 and MSA180, as well as the autocatalytic processing site within SERA6.

In previous work we showed that the MSA180 p40 and p45 fragments that interact with SERA6 PAP are derived from segments of MSA180 that are non-contiguous in the primary sequence and that formation of these fragments results from at least one internal cleavage mediated by SUB1 [35, 23]. However, there is no experimental structural information on MSA180. To examine this issue in more detail we performed structural modelling of MSA180 using AlphaFold3. Intriguingly, this revealed that p40 and p45 are predicted to exist as folded domains that interact closely within the intact FL MSA180 (Fig 1 and Supplementary Figure S1). Moreover, the C-terminal region of MSA180 p45 is predicted to possess a fold belonging to the ubiquitin-like (UBL) family of proteins [36] that is solvent exposed and conserved across orthologues of MSA180 in other *Plasmodium* species. The significance of this was not further explored in this study, but it is conceivable that the MSA180 p45 UBL domain could play a role in its interactions with SERA6 PAP and/or component(s) of the RBC cytoskeleton.

### Mature SERA6 is an enzymatically active protease

SERA6 maturation (conversion to PAP) in intraerythrocytic parasites is temporally associated with cleavage of host RBC β-spectrin; this is thought to lead to RBC membrane rupture through breakdown of the RBC cytoskeleton [13, 23]. The major cleavage site within β-spectrin was mapped to a site close to its N-terminus, between the tandem calponin homology (CH) domains of the actin-binding region of β-spectrin. No amino acid sequence resembling this β-spectrin cleavage site is identifiable within the SERA6 sequence, but since prodomain cleavage during conversion of SERA6 p65 to PAP appears to be autocatalytic (as is common for many papain-like cysteine proteases [37]), we reasoned that small peptides based on the internal cleavage site(s) within SERA6 p65 might act as substrates for PAP *in trans*. However, the internal cleavage site within SERA6 p65 has not been experimentally identified.

In cysteine proteases of the papain-like family, the S2 pocket of the active site cleft (Schechter and Berger nomenclature: [38]) generally primarily defines enzyme specificity, with a common preference for hydrophobic and aromatic residues at the interacting P2 position of substrate [39]. In the AlphaFold2 model of SERA6, a Tyr residue (Y783) forms the floor of the hydrophobic S2 pocket, restricting its size to favour interaction with Leu over larger hydrophobic residues. Similar to proprotease forms of two of the most closely related human cysteine proteases, cathepsin B (CatB) [40] and cathepsin L (CatL) [41, 42], the SERA6 AlphaFold2 model indicates a portion of the prodomain interacting with and obscuring the enzyme active site in a reverse orientation to substrate (thus preventing cleavage), with a Leu sidechain (L489) filling the S2 pocket in SERA6. Autocatalytic activation of the cathepsins involves both unimolecular and bimolecular (or intermolecular) processes [43–45] and in each case prodomain cleavage to generate the mature enzyme occurs at a position downstream from the site that initially fills the active site. At the primary sequence level these cleavage sites in CatB and CatL show little similarity to SERA6 (Fig 4A). To attempt to identify the putative internal SERA6 autocatalytic cleavage site, we therefore compared the AlphaFold2 model of *P. falciparum* SERA6 with the x-ray crystal structures of CatB and CatL (Fig 4B), focusing on the sites at which prodomain cleavage occurs in the cathepsins. Together with multiple sequence alignment of the SERA6 orthologues from several malaria parasite species, this analysis allowed us to tentatively identify a putative prodomain cleavage site within *P. falciparum* SERA6 (Fig 4A) that is conserved across SERA6 orthologues and includes Leu (L604 in *P. falciparum* SERA6) at the P2 position.

**Figure 4.**
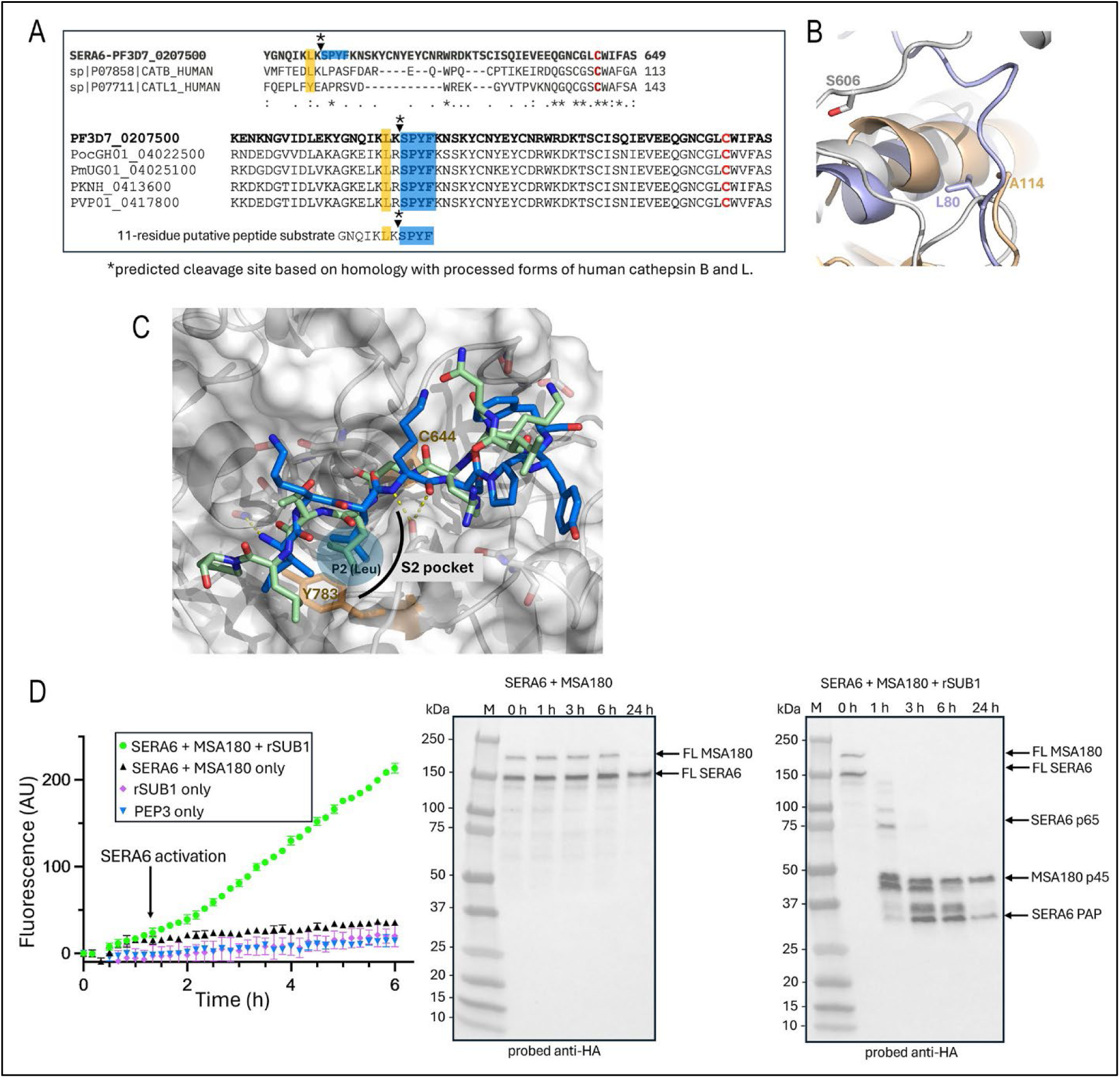
Putative identification of the SERA6 autocatalytic cleavage site. (A) Top, ClustalW primary sequence alignment of partial sequences of *P. falciparum* 3D7 SERA6 (PF3D7_0207500) with human CatB (UniProt P07858) and CatL (UniProt P07711). The prodomain cleavage sites in human CatB and CatL are indicated (arrowhead and an asterisk), with the preceding P2 residue highlighted in mustard. The catalytic Cys residue in each case (C644 in *P. falciparum* SERA6) is in red. Bottom, ClustalW primary sequence alignment of partial sequences of SERA6 orthologues from *P. falciparum*, *P. ovale* (PocGH01_04022500), *P. malariae* (PmUG01_04025100) *P. knowlesi* (PKNH_0413600) and *P. vivax* (PVP01_0417800). The predicted SERA6 autocatalytic cleavage site, based on structural comparisons between the SERA6 AlphaFold models and human CatL and CatB is indicated by an asterisk and arrowhead above the alignment. The conserved P1’-P4’ segment immediately following the predicted scissile bond is highlighted in blue, whilst the conserved catalytic Cys is in red. The sequence of peptide GNQIKLKSPYF which contains the predicted auto-processing site is shown below the alignment with the conserved prime region again highlighted in blue. (B) Cartoon representation of superimposed SERA6 AlphaFold2 model (Q9TY96) in grey, CatB (PDB: 3PBH) in light purple and CatL (PDB: 1MHW) in gold. The known (L80 for CatB and A114 for CatL) and predicted (S606 for SERA6) processing sites are illustrated by their side chain residues. These residues are in vicinity of each other and located in flexible loops near the active site. (C) Zoomed view of molecular representation of the SERA6 AlphaFold2 model shown as a light grey cartoon with a semi-transparent molecular surface. Side chains of the catalytic C644, as well as residue Y783 which is predicted to lie at the bottom of the S2 pocket, are shown as orange sticks. The SERA6 prodomain segment that lies in the SERA6 active site in an opposite orientation to substrate is shown as green sticks coloured by elements (oxygen, red; nitrogen, blue). A Leu residue is found in the S2 pocket. The docked GNQIKLKSPYF peptide (blue) (ICM-Pro score -20) is shown superimposed in a substrate-binding orientation, showing its P2 Leu also in the SERA6 S2 pocket. (D) Left hand side, progress curve showing fluorescence increases upon cleavage of fluorescent peptide substrate PEP3 in a typical assay containing affinity-purified FL SERA6 and MSA180 with and without addition of rSUB1. Minimal fluorescence increases were observed in the control reactions. Right, western blot analysis of supernatants from the fluorescence assays containing FL SERA6 and FL MSA180, with and without rSUB1. The appearance of detectable SERA6 PAP approximately correlates with the initiation of fluorescence increases.

To test this prediction, we first took an 11-residue peptide (GNQIKLKSPYF) corresponding to that spanning the putative prodomain cleavage site in SERA6 and used *in silico* docking into the AlphaFold2 model of the SERA6 papain-like catalytic domain to assess the capacity of the peptide to be accommodated in a substrate-like manner in the SERA6 active site. As shown in Fig 4B, the peptide docked well into the SERA6 model, with the conserved L604 sidechain interacting with Tyr783 at the bottom of the SERA6 S2 pocket. Encouraged by this, we next produced a quenched fluorogenic peptide (Dabcyl-GNQIKLKSPYF-EDANS, called PEP3) based on the 11-mer peptide and designed such that proteolytic cleavage anywhere within the peptide backbone should produce a fluorescence increase. To assess the capacity of PEP3 to be cleaved by active SERA6, we added PEP3 to assays similar to those described in Fig 3D, containing reconstituted FL SERA6 and FL MSA180 affinity-purified from parasite saponin extracts. Increases in fluorescence corresponding to cleavage of PEP3 occurred following addition of rSUB1 (Fig 4C). Interestingly, the fluorescence increase exhibited a kinetic profile that encompassed a time delay or lag phase that approximately correlated with the period before the detectable appearance of SERA6 PAP, consistent with the notion that conversion to PAP was required for the initiation of PEP3 cleavage. Analysis of the assay supernatants by reversed-phase HPLC (RP-HPLC) and electrospray mass spectrometry to detect the fluorescent (EDANS-containing) product(s) of PEP3 cleavage revealed that PEP3 was cleaved in these assays primarily or exclusively at the Lys-Ser bond corresponding to the predicted autocatalytic cleavage site within SERA6 at the IKLK-SPYF bond (hyphen indicating the scissile bond) (Fig 4A and Supplementary Fig S2). Importantly, no PEP3 cleavage was detected in control incubations containing rSUB1 only, in accord with the well-established substrate specificity of SUB1 [32, 46] which is not predicted to be capable of cleaving PEP3 anywhere within its sequence. It was concluded that maturation of SERA6 to PAP resulted in the generation of a proteolytic activity capable of cleaving PEP3 at a preferred internal site corresponding precisely to the predicted autocatalytic site within SERA6.

### Development of improved drug-like inhibitors of SERA6

We have previously described [23] two structurally-related nitrile compounds that, similar to the epoxide E64 and its more membrane-permeable analogue E64-d, inhibit both the final autocatalytic step of SERA6 maturation and SERA6-mediated cleavage of β-spectrin in *P. falciparum* schizonts undergoing egress. Both these nitrile compounds, a purine called MMV676881 and the triazine compound 31, also block parasite egress with a phenotype typical of inhibition or genetic ablation of SERA6 [23, 29]. This is characterised by normal rounding and rupture of the PVM with the characteristic increased mobility of the intraerythrocytic merozoites, concomitant with poration of the RBC membrane but failure of the RBC membrane to break [13]. The two nitrile compounds were similarly potent (EC_50_ ∼0.4 μM for inhibition of SERA6 maturation in intraerythrocytic parasites) but were predicted to be poorly soluble. To seek analogues with improved potency and/or solubility, we screened a small set of 30 related nitrile compounds (a kind gift of Min Shen and Matthew Hall, National Center for Advancing Translational Sciences [NCATS], NIH, USA) for their capacity to inhibit egress, using a previously-described assay based on spectrometric detection of haemoglobin released from live schizonts upon egress [23]. This screen identified two further active (defined as an egress-inhibitory EC_50_ <2 μM) purine nitrile compounds called compounds 33 and 37 (Fig 5A). Further examination showed that the newly identified hits were similarly potent as the previously-identified MMV676881 and compound 31 (which were also originally derived from the same NCATS set) in inhibition of SERA6 p65-to-PAP conversion in intact *SERA6-mTAP:loxP* schizonts (Fig 5B), confirming the new compounds as inhibitors of SERA6 maturation. Notably, all four active compounds shared a 3,5-difluorophenyl moiety. Comparison with the inactive nitriles in the NCATS screening set suggested that replacement or modification of this 3,5-difluorophenyl group results in loss of inhibitory activity (Supplementary Fig S3). We also noted that cyclisation of the triazine compound 31 in two different vectors could rationalise the different architecture of the three purine nitrile actives (Supplementary Fig S3). All the NCATS nitrile compounds were originally described in an earlier study by Mott and colleagues [47] in which MMV676881 (referred in that work as compound 32) was co-crystallised with the *Trypanosoma cruzi* cysteine protease cruzain. This x-ray crystal structure showed that the cyano group of MMV676881 formed the expected covalent adduct with the nucleophilic active-site Cys of cruzain and the 3,5-difluorophenyl substituent extended into the largely hydrophobic specificity-determining S2 pocket of cruzain, whilst the purine core of MMV676881 orientated the vectors of these two key interacting groups (Fig 5C and Supplementary Fig S4). This binding pose is remarkably similar to that adopted by a number of similar purine nitriles bound to the active site of the papain-like cysteine protease cathepsin K [48]. To examine the possibility that MMV676881 and the other actives might interact in a similar manner with SERA6, we performed *in silico* covalent docking of MMV676881 into the ICM-Pro molecular model of the SERA6 papain-like catalytic domain, comparing this model with the cruzain-MMV676881 and cathepsin K-purine nitrile structures (Fig 5D). This showed that indeed MMV676881 and the three other active compounds described here might interact with SERA6 in a very similar manner to the MMV676881-cruzain and the cathepsin K-nitrile compound interactions.

**Figure 5.**
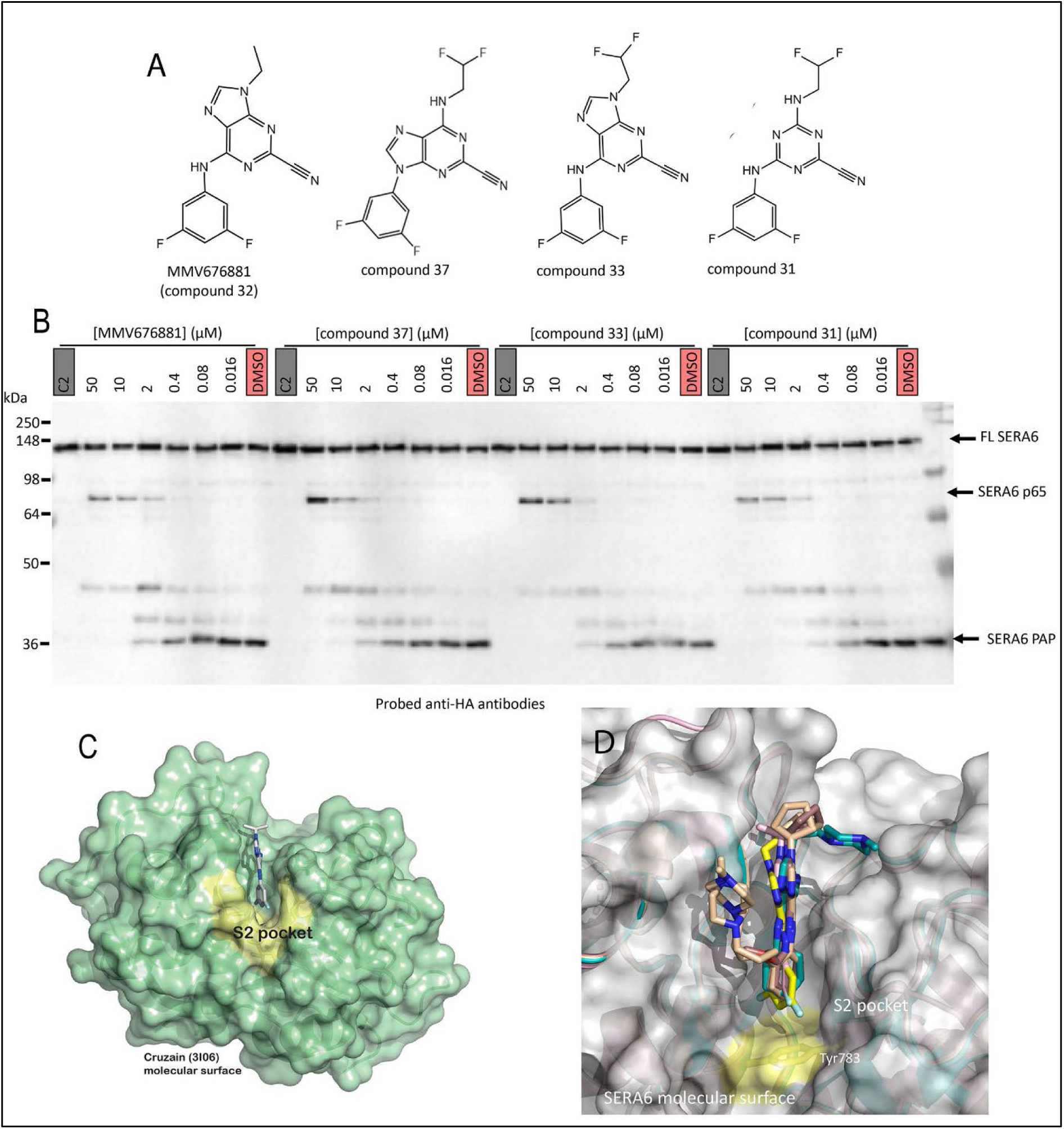
New purine nitrile inhibitors of SERA6 maturation. (A) Structures of the newly identified egress inhibitors compounds 33 and 37, alongside the previously-identified hits MMV676881 and the triazine compound 31. (B) Western blot-based assay showing dose-dependent inhibition of SERA6 maturation (SERA6 PAP formation) in intact *SERA6-mTAP:loxP* schizonts treated with the indicated concentrations of the triazine and purine nitrile hits. Vehicle only (DMSO) was used as a negative control in all assays. C2 is an inhibitor of PKG-mediated SUB1 discharge, so blocked all SERA6 maturation in the schizonts. (C) Model of the x-ray crystal structure (PDB: 3I06) of MMV676881 bound to the *Trypanosoma cruzi* papain-like cysteine protease cruzain [47]. Cruzain is shown as a semi-transparent green molecular surface overlaying a ribbon diagram, with the specificity-determining S2 pocket highlighted in yellow. (D) *In silico* covalent docking of MMV676881 (yellow sticks coloured by elements oxygen, red and nitrogen, blue) into the AlphaFold2 molecular model (grey molecular surface) of the SERA6 papain-like catalytic domain. Note the predicted Y783 residue at the base of the SERA6 S2 pocket. The model is shown superimposed onto the x-ray crystal stuctures of cruzain bound to MMV676881 (pink, PDB: 3I06, rmsd deviation from the MMV676881 model 0.919 Å); cathepsin K bound to purine nitrile NVP-ABE854 (light blue, PDB: 1U9V, rmsd 0.930 Å); cathepsin K bound to NVP-ABI491 (dark brown, PDB: 1U9W, rmsd 0.935 Å); and cathepsin K bound to NVP-ABJ688 (light brown, PDB: 1U9X, rmsd 0.929 Å).

Based on our structural predictions of the MMV676881-SERA6 interaction and pharmacophore analysis of the active compounds, we designed an additional series of eight analogues (all named with the prefix CS) that we predicted to be more soluble and drug-like than the existing four hits, whilst retaining the structural characteristics associated with SERA6 inhibition (see Supplementary Fig S5 and Supplementary Fig S6 for their structures and predicted chemical properties). The new CS compounds were assessed for their capacity to prevent SERA6-mediated β-spectrin cleavage in treated parasites, to inhibit SERA6 maturation in the saponin extract assay developed above, and to prevent parasite replication in *in vitro* culture. As shown in Fig 6A-C, five of the CS compounds showed inhibitory activity in all three assays, with CS01-03 displaying the highest potency in the quantitative cell-based replication assay (which measures the transition from mature schizonts to newly invaded ‘ring’ stage parasites, a surrogate measure of successful egress of invasive merozoites). Notably, in this last assay CS01-03 was ∼4.4-fold more potent than the original hit compound MMV676881. *In silico* covalent docking analysis of the active CS compounds was in agreement with the experimental findings, suggesting that the active CS compounds have the capacity to form additional main-chain H-bond interactions with the SERA6 S2 pocket compared to MMV676881 (Fig 6D). In keeping with its relative potency in the other assays, compound CS01-03 consistently gave the best docking scores in this analysis (Supplementary Fig S7)

**Figure 6.**
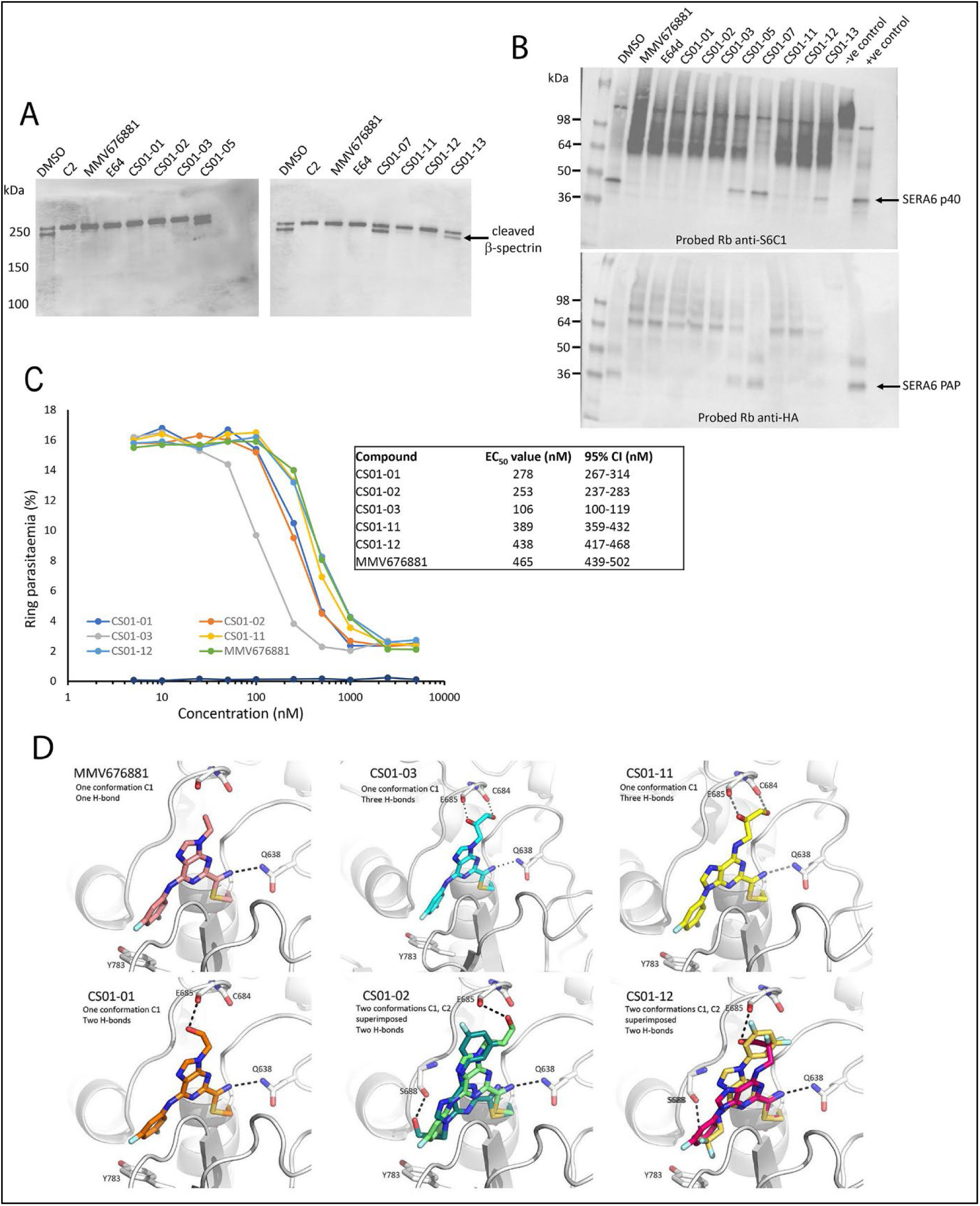
CS01-03 is a drug-like nitrile inhibitor with improved potency against SERA6. (A) Inhibition of β-spectrin cleavage at egress. Western blot-based assay showing the effects on β-spectrin cleavage upon treatment of mature schizonts with the indicated compounds (10 μM final concentration). Compounds CS01-01, CS01-02, CS01-03, CS01-11 and CS01-12 inhibited β-spectrin cleavage in this assay. The PKG inhibitor C2 (which blocks SUB1 discharge) and the known SERA6 inhibitors MMV676881 and E64 were used as positive control inhibitors. Vehicle only (DMSO) acted as a negative control. (B) Inhibition of SERA6 maturation *ex vivo* in schizont saponin extracts. Western blot-based assay showing the effects on SERA6 maturation following addition of rSUB1 to saponin extracts of mature *SERA6-mTAP:loxP* schizonts supplemented with the test CS compounds (10 μM final). Control compounds were as in (A) except that the membrane-permeable E64 analogue E64-d was used in this case. In this experiment the negative control contained no rSUB1, whilst the positive control contained rSUB1 but lacked inhibitors. SERA6 PAP was detected with rabbit anti-HA antibodies whilst the anti-SERA6 antibody anti-S6C1 [22] was used to detect the SERA6 prodomain p40 which is generated upon formation of SERA6 PAP. (C) Typical dose-response plot (of 3 independent assays) showing the effects of the CS compounds and MMV676881 on schizont-to-ring transition. Ring parasitaemia (Y-axis) is shown plotted against drug concentrations ranging from 5 nM to 5 mM (X-axis). Parasitaemia values were determined by flow cytometry. CS01-03 was ∼4.4-fold more potent than the original hit compound MMV676881, with an EC_50_ value of 106 nM. (D) *In silico* covalent docking of the hit purine nitriles into the active site S2 pocket of the AlphaFold2 SERA6 PAP model. CS01-02 and CS01-12 are shown docked in 2 alternative conformations. Note that all the CS actives showed the potential to form additional H-bonds compared to the original active MMV676881. Each compound and key interacting residues of SERA6 are shown as sticks coloured by elements (oxygen, red; nitrogen, blue). All analysis was performed in ICM-Pro.

In view of these promising results, we next focused on CS01-03. This compound possesses a chiral carbon atom and further docking studies suggested that the *R*-enantiomer likely binds to SERA6 better than the *S*-enantiomer (Supplementary Fig S8). To examine this experimentally, the CS01-03 racemate was subjected to chiral fractionation, resulting in the isolation of the two enantiomers CRK1_1 and CRK1_2. Of these, CRK1_1 was the more potent in the schizont-to-ring transition assay, with an estimated EC_50_ ∼100 nM (Supplementary Fig S9). In vitro drug metabolism and pharmacokinetic (DMPK) analysis of CRK1_1 and CRK1_2 indicated good solubility and stability for both enantiomers (Supplementary Fig S10). To determine whether the antimalarial action of CRK1_1 was through selective inhibition of SERA6 maturation as anticipated, CRK1_1 was evaluated in a range of further assays to establish its selectivity and mode of action. In the first of these assays, synchronous cultures of ring-stage parasites treated with CRK1_1 (500 nM, ∼5-fold EC_50_) were monitored by microscopic examination at intervals over the course of an entire ∼48 h erythrocytic cycle. As shown in Fig 7A, whilst some mild swelling of the digestive vacuole was apparent in treated parasites at the early trophozoite (31 h post-invasion) developmental stage (possibly due to inhibition of the non-essential papain-like cysteine protease haemoglobinase falcipain 2 [49]), crucially CRK1_1 treatment did not block parasite development until the mature schizont stage, where complete arrest of egress was observed. This indicated a good degree of selectivity for CRK1_1. In vitro enzyme assays supported this; whilst at high concentrations (10 μM) CRK1_1 showed some activity against mammalian papain, the compound showed no inhibition of rSUB1 or recombinant PMX (Supplementary Fig S11). This confirmed that the effects of CRK1_1 on egress were not due to inhibition of these crucial parasite proteolytic enzymes.

**Figure 7.**
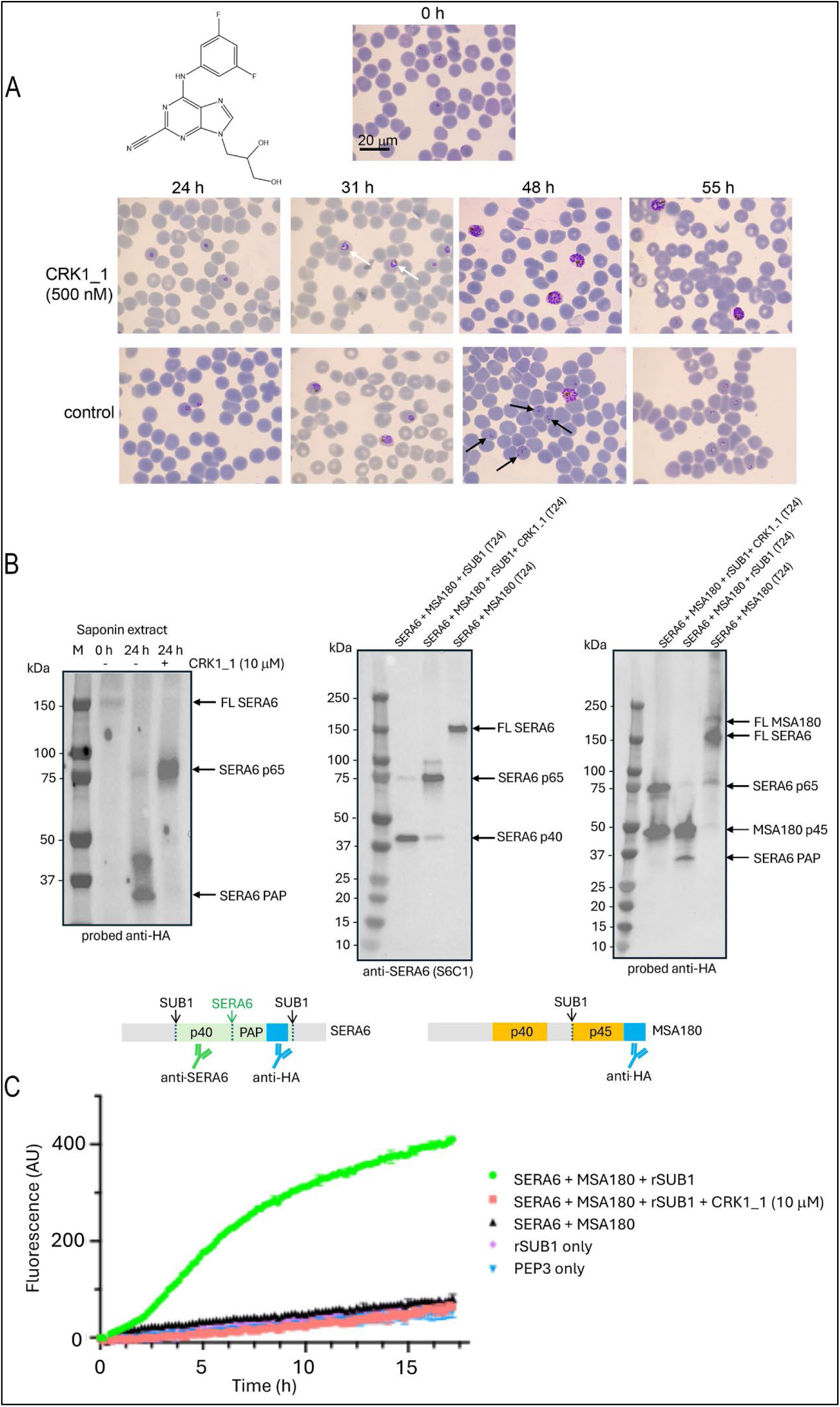
CRK1_1 exerts its antimalarial effects through inhibition of SERA6 maturation. (A) Top left, structure of CRK1_1. The stereochemistry is not indicated, as it was not determined whether CRK1_1 was the *R*- or *S*-enantiomer of CS01_03. Right and below; synchronous *P. falciparum* cultures at early ring stage (0 h) were supplemented with either CRK1_1 (500 nM) or vehicle only (1% [v/v] DMSO, control) then examined microscopically by Giemsa stain at the indicated intervals over the course of the entire ensuing erythrocytic cycle. Some vacuolation (white arrows) was evident in the CRK1_1-treated parasites by 31 h post-invasion but the parasites continued to develop without discernible retardation to mature schizont stage then failed to egress. In contrast, in control cultures multiple new rings (black arrows) were present as expected by 48 h, indicative of successful egress, and all egress was complete by 55 h. (B) CRK1_1 arrests SERA6 maturation at the p65 intermediate stage. Saponin extracts of *SERA6-mTAP:loxP* schizonts (left-hand side) or reconstituted, affinity-purified HA-tagged FL SERA6 and FL MSA180 ( middle and right-hand side ) were incubated with rSUB1 in the presence or absence of CRK1_1 and formation of SERA6 p65, the free prodomain SERA6 p40 and SERA6 PAP assessed by Western blot. CRK1_1 inhibited conversion of SERA6 p65 to the mature PAP and the free prodomain. (C) CRK1_1 inhibits SERA6 peptidase activity. Cleavage of the fluorogenic SERA6 substrate PEP3, as measured by an increase in fluorescence, is inhibited by CRK1_1.

CRK1_1 was finally evaluated in the newly developed *in vitro* reconstitution assay and SERA6 peptidase activity assay described above. As shown in Fig 7B, SUB1-dependent maturation of SERA6 in the presence of CRK1_1 was arrested at the p65 intermediate form, as expected, indicating direct inhibition of the autocatalytic p65-to-PAP step of maturation. Furthermore, CRK1_1 prevented cleavage of the fluorogenic substrate PEP3 in the SERA6 activity assay (Fig 7C and Supplementary Fig S12), showing a direct on-target effect of CRK1_1 on the proteolytic activity of SERA6 PAP. It was concluded that CRK1_1 kills the parasite primarily or exclusively through prevention of egress as a result of inhibition of SERA6 maturation and activity. Importantly, CRK1_1 displays improved potency, physicochemical properties and drug-likeness compared to the original hit MMV676881. CRK1_1 represents the most potent and selective drug-like inhibitor of *P. falciparum* SERA6 activity described to date.

## Discussion

Egress is a crucial step in the lifecycle of the blood-stage malaria parasite, being essential for parasite replication and directly causing much of the pathophysiology of the periodic fevers associated with the disease due to the release into the circulation of large amounts of parasite antigen upon schizont rupture. Whilst no antimalarial drugs currently in clinical use specifically target egress, it has become increasingly clear that egress involves several parasite enzymes that have high potential as targets for new types of chemotherapeutic approaches (for comprehensive recent reviews see [50, 51]. The present study significantly advances our understanding of the maturation of SERA6 and its role in malaria parasite egress. Using unfractionated saponin extracts derived from mature schizonts arrested with an inhibitor of SUB1 discharge (the PKG inhibitor C2), we first showed that simple supplementation of the extracts with rSUB1 was sufficient to obtain complete maturation of the endogenous FL SERA6 to its terminal PAP form. The kinetics of SERA6 maturation in these *in vitro* experiments were relatively slow (requiring several hours compared to the very rapid process in the parasite which takes just minutes; [23]). We do not know the reason for this but we rationalise that it may be a result of suboptimal conditions of salt, pH or reactant concentrations in our assays. Further refinement of these experiments by affinity purification of FL SERA and FL MSA180 and reconstitution of the isolated proteins in the presence of rSUB1, combined with the development of a fluorogenic substrate based on a putative internal SERA6 cleavage site, allowed us to show that SUB1 and MSA180 alone are necessary and sufficient for maturation of SERA6 to an active protease (more accurately, peptidase). Our experimental approaches cannot formally rule out the possibility that other essential parasite components co-purified with either FL SERA6 or FL MSA180 in our affinity purification procedures, but we consider this unlikely in the lack of any evidence that the precursor proteins form heterocomplexes in the parasite PV.

Our study reinforces the essentiality of MSA180 for SERA6 maturation, but the molecular function of MSA180 remains unknown. We have previously speculated [23] that MSA180 may provide a scaffolding or chaperone-like role during SERA6 maturation and/or the presumed RBC cytoskeletal interaction required for β-spectrin cleavage. Intriguingly, our bioinformatic analysis using AlphaFold3 has shown that the p45 region of *P. falciparum* MSA180 (one of the two fragments of MSA180 that binds to SERA6 PAP) contains a solvent-exposed UBL domain that is predicted to be conserved across MSA180 orthologues in other *Plasmodium* species. UBL domains are often involved in protein-protein interactions [52, 53]. The MSA180 UBL domain is predicted to be less solvent-exposed within the fold of the FL MSA180 prior to its cleavage to generate the p40 and p45 domains, so it is tempting to speculate that MSA180 processing may potentially function to expose the UBL element to enable it to partake in binding interactions with SERA6 PAP or indeed with β-spectrin or other associated components of the RBC cytoskeleton. Remarkably, the AlphaFold3 predictions also support a stable pre-existing non-covalent interaction between the p45 and p40 segments within the intact FL MSA180, consistent with our evidence that both fragments interact with SERA6 PAP [23]. Further experimental work will be required to establish the precise role(s) of MSA180 and its UBL domain in SERA6 maturation and activity.

Our demonstration of the appearance of peptidase activity associated with maturation of FL SERA6 or p65 to PAP constitutes the first experimental evidence to our knowledge that SERA6 PAP (but not the precursor FL SERA6) is an active proteolytic enzyme. This is consistent with the previously established temporal association between maturation of SERA6 and cleavage of host RBC β-spectrin at egress. Our use of saponin extracts as a source of materials for analysing SERA6 maturation provides a simple but significant technical advance. Previously, screening for inhibitors of maturation has required work with continuous cultures of live *P. falciparum* asexual blood-stage parasites, with all the associated need for high containment culture facilities, laborious parasite manipulation approaches and ready access to human blood. Because saponin extracts can be generated in bulk and frozen in aliquots, it is now feasible to disseminate extracts to multiple laboratories if required to enable their use for multiple downstream assays under non containment conditions. Our development of a simple, cell-free assay that measures SERA6 PAP activity using a synthetic fluorogenic peptide substrate will greatly accelerate the development and optimisation of new selective small molecule inhibitors of SERA6. Our development of improved small molecule inhibitors of SERA6 activity and egress in this study validates the SERA6 activity assay.

CRK1_1, the more active enantiomer of CS01-03, is the most potent and selective small molecule inhibitor of SERA6 maturation identified to date. Orally available carbonitrile compounds have been used in the clinic for some time (e.g. as inhibitors of dipeptidyl peptidase-IV and the SARS-CoV-2 main protease; [54, 55]) so further development of CRK1_1 or its analogues as a drug lead or experimental tool is warranted. Our detailed phenotypic analysis of parasites treated for an entire erythrocytic lifecycle with CRK1_1 indicated swelling of the parasite food vacuole, typical of compounds that inhibit the haemoglobin digestion pathway. This was not unexpected, since haemoglobin catabolism involves the activity of parasite falcipains [56], a class of four *P. falciparum* papain-like cysteine proteases, one or more of which is likely susceptible to inhibition by CRK1_1 due to their structural relatedness to SERA6. Our observation that this cross-inhibition was not sufficiently deleterious to arrest parasite growth was unsurprising, as extensive work by others has concluded that proteases within the haemoglobin digestion pathway are highly redundant, to the extent that several are no longer considered good targets for antimalarial drug development [56, 57]. On the other hand, at least one falcipain appears to play crucial role(s) during the asexual blood stage life cycle [56]. Compounds that kill the parasite through the inhibition of multiple targets are particularly attractive for drug development as they are generally expected to be less likely to select for parasite resistance than drugs that target a single pathway. To this end it may be possible to identify CRK1_1 analogues that potently inhibit SERA6 maturation as well as important falcipain functions, placing additional pressure on parasite survival and reducing opportunities for parasite resistance.

## Acknowledgements

The authors are indebted to the Medicines for Malaria Venture (MMV) for provision of compound MMV676881. This work was also supported by funding to MJB from the Francis Crick Institute (https://www.crick.ac.uk/), which receives its core funding from Cancer Research UK (FC001043; https://www.cancerresearchuk.org), the UK Medical Research Council (FC001043; https://www.mrc.ac.uk/), and the Wellcome Trust (FC001043; https://wellcome.ac.uk/). M.S.Y.T was in receipt of a Francis Crick PhD studentship, as well as funding from the Francis Crick Idea to Innovation (i2i) programme (grant P2019-0015). The work was also supported by Wellcome ISSF2 funding to the London School of Hygiene & Tropical Medicine, and the authors are immensely grateful to Min Shen and Matthew Hall at the intramural research program of the National Center for Advancing Translational Sciences (NCATS), Division of Pre-Clinical Innovation, USA, for the valuable gift of the purine and triazine nitrile library. We thank Reach Separations (Bio City, Pennyfoot St, Nottingham, UK) for the chiral fractionation of the CS01-03 racemate, and we are grateful to Jennifer Riley (Dundee Drug Discovery Unit) for the in vitro DMPK analysis of CRK1_1 and CRK1_2. For the purpose of Open Access, the author has applied a CC BY public copyright licence to any Author Accepted Manuscript version arising from this submission.

## Materials and Methods

### Transgenic *P. falciparum* lines and maintenance

*P. falciparum* 3D7 and the transgenic *P. falciparum* lines *SERA6-mTAP:loxP* and *MSA180-HA3:loxP* were generated and maintained *in vitro* as described previously [23]. To generate saponin extracts, highly mature schizonts enriched from the synchronised lines by arresting egress with C2 [21] were pelleted and resuspended at room temperature in a buffer composed of Tris-buffered isotonic saline (TBS) supplemented with 2 mM DTT, 2 µM C2 and 0.15% (w/v) saponin. The resulting extracts, enriched in released PV proteins, were clarified by centrifugation, snap frozen and stored in aliquots at -70°C until required.

### Compounds and peptides

The CS compounds were synthesised by Enamine Ltd (Kyiv, Ukraine). Compound MMV676881 (a kind gift of MMV), the PKG inhibitor C2 (a kind gift of Simon Osborne, LifeArc UK), the NCATS compounds and the CS compounds were stored at -20°C in dry 100% DMSO at a concentration of 2-10 mM. Fluorogenic peptide PEP3 (Dabcyl-GNQIKLKSPYF-EDANS) and the HA peptide used for elution of HA-tagged protein from anti-HA antibody beads were synthesised in-house by the Crick Chemical Biology Science Technology Platform and were stored as 1 mM stock solutions in DMSO at -20°C.

### Affinity purification of HA-tagged parasite SERA6 and MSA180

Saponin extracts from C2-treated schizonts of the *P. falciparum SERA6-mTAP:loxP* and *MSA180-HA3:loxP* transgenic lines were processed in parallel using anti-HA antibody magnetic beads (Pierce) according to the manufacturer’s instructions (Thermo Fisher Scientific). Briefly, 50 µl aliquots of a suspension of magnetic beads were washed twice in TBS, 25 mM CHAPS, 2 mM DTT while aliquots (400-800 μl) of frozen saponin extracts containing HA-tagged SERA6 and MSA180 in TBS buffer, 2 mM DTT, 2 μM C2 and 0.15% (w/v) saponin with no additional protease inhibitors were thawed on ice. The extracts were supplemented with CHAPS to 25 mM final before being added to the washed magnetic beads. The resulting suspension was incubated on a rotating wheel for 2 h at 4°C to allow binding. Unbound material was then removed and the beads washed five times in TBS, 25 mM CHAPS, 2 mM DTT before eluting bound protein by incubation with 200 µl of a solution of HA peptide (2 mg/ml in TBS) for 2 h at 37°C. The eluted purified SERA6 and MSA180 were usually immediately used for reconstitution/processing and activity assays, or alternatively were stored at 4°C for a maximum of 24 h before use. The starting saponin extracts, the unbound material, the first wash, the eluate samples and the beads (before and after elution) were retained for analysis by SDS PAGE and western blotting using nitrocellulose membranes (GE Healthcare). Membranes were probed with the primary anti-HA rat monoclonal antibody 3F10 (diluted 1:2,000), or a previously-described rabbit polyclonal anti-SERA6 antibody (called anti-S6C1: [22]) (1:1,000), or a polyclonal rabbit anti-MSA180 antibody (1:500) [23]. HRP-conjugated polyclonal goat anti-rat (1:10,000) or goat anti-rabbit (1:3,000) antibody were used as secondary antibodies, followed by detection using Immobilon Western Chemiluminescent HRP Substrate (Millipore) according to the manufacturer’s instructions. Blots were visualised and documented using a ChemiDoc Imager (Bio-Rad) with Image Lab software (Bio-Rad).

### Saponin extract, SERA6 and MSA180 reconstitution assays and SERA6 peptidase activity

Whole schizont saponin extracts or eluted, affinity-purified FL SERA6 and FL MSA180 proteins were combined as required (usually 1:1 for the purified proteins) and made to 25 mM CHAPS, 15 mM CaCl_2_ (a requirement for SUB1 activity) before adding 1.71 U/µl of rSUB1 in 50 mM Tris-HCl, pH 8.0, 150 mM NaCl. Processing was allowed to occur overnight at room temperature or 37°C, then assessed by western blotting analysis.

SERA6 peptidase activity was assessed in white 96-well opaque flat-bottom plates (Nunc) by monitoring fluorescence increases upon cleavage of the fluorogenic peptide PEP3 (Dabcyl-GNQIKLKSPYF-EDANS), used at final concentration of 10 µM in the assay. For each condition, a final volume of 200 µl/well was used with the following reaction buffer: 1xTBS, 25 mM CHAPS, 15 mM CaCl_2_, 6 mM DTT and 1% DMSO. Fluorescence changes were monitored overnight at room temperature in a SpectraMax M5e Multi-Mode Microplate Reader (Molecular Devices LLC) (Ex 340 nm, Em 492 nm), taking readings at intervals of 10 min and blanking against reaction buffer only. Assays were carried out in duplicate and GraphPad Prism software was used to analyse the data. At the end of each run, the protein solutions from the flat-bottomed plates were extracted directly into SDS and stored at -20°C prior to SDS-PAGE and western blot analysis.

### CRK1_1 selectivity assays

Compound CRK1_1 was assessed for inhibitory activity against rSUB1, PMX and papain. Purified rSUB1 was diluted 1:500 from a stock solution (228 U/ml stock in 20 mM Tris–HCl pH 8.2, 150 mM NaCl, 10% glycerol) into SUB1 reaction buffer (20 mM Tris–HCl pH 8.2, 150 mM NaCl, 12 mM CaCl_2_, 25 mM CHAPS). This was supplemented with CRK1_1 from a 100% DMSO stock solution (1 mM stock, 10 μM final). Following a 5 min incubation, the mixture was supplemented with the fluorogenic peptide SUB1 substrate SERA4st1F-6R12 [46] (Ac-CKITAQDDEESC-OH labelled on both Cys side-chains with tetramethylrhodamine; 0.1 μM final). Recombinant PMX [58] was similarly used at a final concentration of 0.36 μM in 25 mM sodium acetate pH 5.5, 0.005% Tween 20 reaction buffer, in this case using fluorogenic substrate Dabcyl-HSFIQEGKEE-EDANS (Ex 340 nm, Em 492) at a final concentration of 1.6 μM in the reaction buffer. The potent PMX inhibitor 7k [58, 59] was used as a control (10 μM final). Commercially-obtained papain from papaya latex (Sigma, P4762) was diluted to 0.1 mg/ml in 25 mM Bis-Tris pH 6.5, 1 mM EDTA, 5 mM DTT and incubated for 30 min in the reaction buffer to allow activation before use at 0.01 mg/ml. In this case fluorogenic substrate Nα-Benzoyl-DLR-AMC (1mM stock in 100% DMSO) (Ex 380 nm, Em 460 nm) was used, at 10 μM final. The inhibitor control for papain was E64 (10 μM final). All assays were performed in white 96-well opaque flat-bottom plates (Nunc) at room temperature in a SpectraMax M5e Multi-Mode Microplate Reader (Molecular Devices LLC), taking readings at 3 min intervals and blanking with the reaction buffer. All assays were performed in duplicate and GraphPad Prism software used to analyse the data.

### PEP3 cleavage mapping by reversed phase chromatography and mass spectrometry

Reversed-phase high pressure liquid chromatography (RP-HPLC) was used to monitor PEP3 cleavage. Samples (50 μl) from assays containing purified FL SERA6 and FL MSA180 treated with rSUB1 in the presence of fluorogenic peptide PEP3 as described were injected onto a Vydac C18 RP-HPLC column (4.6 mm x 25 cm) using a 100 μl loop and eluted with a 0-90% (v/v) gradient of acetonitrile in 0.1% (v/v) trifluoroacetic acid over a 30 min period, monitoring both absorbance (at 215 nm) and fluorescence (Ex 340 nm, Em 492 nm). Peaks of interest were collected manually and subjected to LC-MS analysis to identify their makeup. Briefly, a M-Class LC (Waters, UK) delivered a flow of approximately 50 µl/min. A C4 UPLC BEH 1.7µm, 2.1 x 5 mm pre-column (Waters, UK), trapped the injected peptides prior to separation on a C4 BEH 1.7 µm, 1.0 x 100 mm UPLC column, eluting with a 20-min gradient of acetonitrile (2% to 80%). The analytical column outlet was directly interfaced via an electrospray ionisation source, with a hybrid quadrupole time-of-flight (Q-TOF) mass spectrometer (SynaptG2Si, Waters, UK). Data was acquired over an *m/z* range of 350–4000, in positive ion mode with a cone voltage of 40 v. Scans were summed together manually and deconvoluted using MaxEnt1 (Masslynx, Waters, UK).

### SERA6 homology modelling, covalent and noncovalent docking of nitrile compounds and peptides

SERA6 homology modelling was performed in ICM-Pro (version3.9-1c/MacOSX, Molsoft LLC, San Diego, CA) and based on the SERA5 proenzyme template (PDB: 6X44, 2.17Å), and using the AlphaFold2 model (Q9TY96), the ‘catalytic pocket’ of which was in an active induced binding mode to accommodate a portion of the inhibitory prodomain. The ICM Full Model Builder procedure was used, making use of the ICM force field and resulting in a fully refined model (including side chains and loops) with a good stereochemistry validation (1.04% outliers). Flexible covalent docking of the nitrile compounds into the active site of the SERA6 model was also performed in ICM-Pro. Protonation of the predicted catalytic histidine H810 (Nδ1) was enabled to favour inhibitor stabilization via hydrogen-bonding with the thiolate form of the catalytic cysteine C644. The SERA6 prodomain binding region (N485-P492) was initially used to define the boundaries within the enzyme active site pocket for the docking procedure, then the entire prodomain region (D371-N611) was removed prior to docking. Potential energy maps of the SERA6 receptor and docking preferences were set up using the program default parameters. Energy terms were based on the all-atom vacuum force field ECEPP/3 and conformational sampling was based on the biased probability Monte Carlo (BPMC) procedure. The nitrile-based CS compounds were derived from the original purine and triazine nitrile actives and were rationally designed using the ICM-Pro ligedit option. Three independent docking runs were performed, with a thoroughness (length of simulation) varied from 3 to 10 and the selection of 3 docking poses. Ligands were ranked according to their ICM energetics (ICM score, unitless), which weighs the internal force-field energy of the ligand combined with other ligand-receptor energy parameters.

Flexible non-covalent docking of the SERA6-derived peptide (I602-F609 of the SERA6 primary amino acid sequence) was also performed using ICM-Pro. As before, the SERA6 prodomain segment (N485-P492) was used to define the active site SERA6 receptor pocket and then removed from the receptor. Potential energy maps of the SERA6 receptor pocket and docking preferences were set up using the program default parameters. Three independent docking runs were performed, with a thoroughness (length of simulation) varying from 5 to 10 and the selection of 2 docking poses.

### Structural modelling of MSA180

MSA180 was modelled using the AlphaFold3 server (https://alphafoldserver.com) using the MSA180_PF3D7_1014100 amino acid sequence as entry. The MSA180 AlphaFold3 model 4 was used for structural investigation with the DALI protein structure comparison server (http://ekhidna2.biocenter.helsinki.fi/dali/) to search for similar folds. This identified the ubiquitin-like modifier FAT10 (PDB: 6GF1) as a match for the C-terminal region of the MSA180 p45 domain.

