## Supplemental figures for "Ex vivo maturation of the malaria parasite egress protease SERA6 aids pathway dissection and inhibitor development"

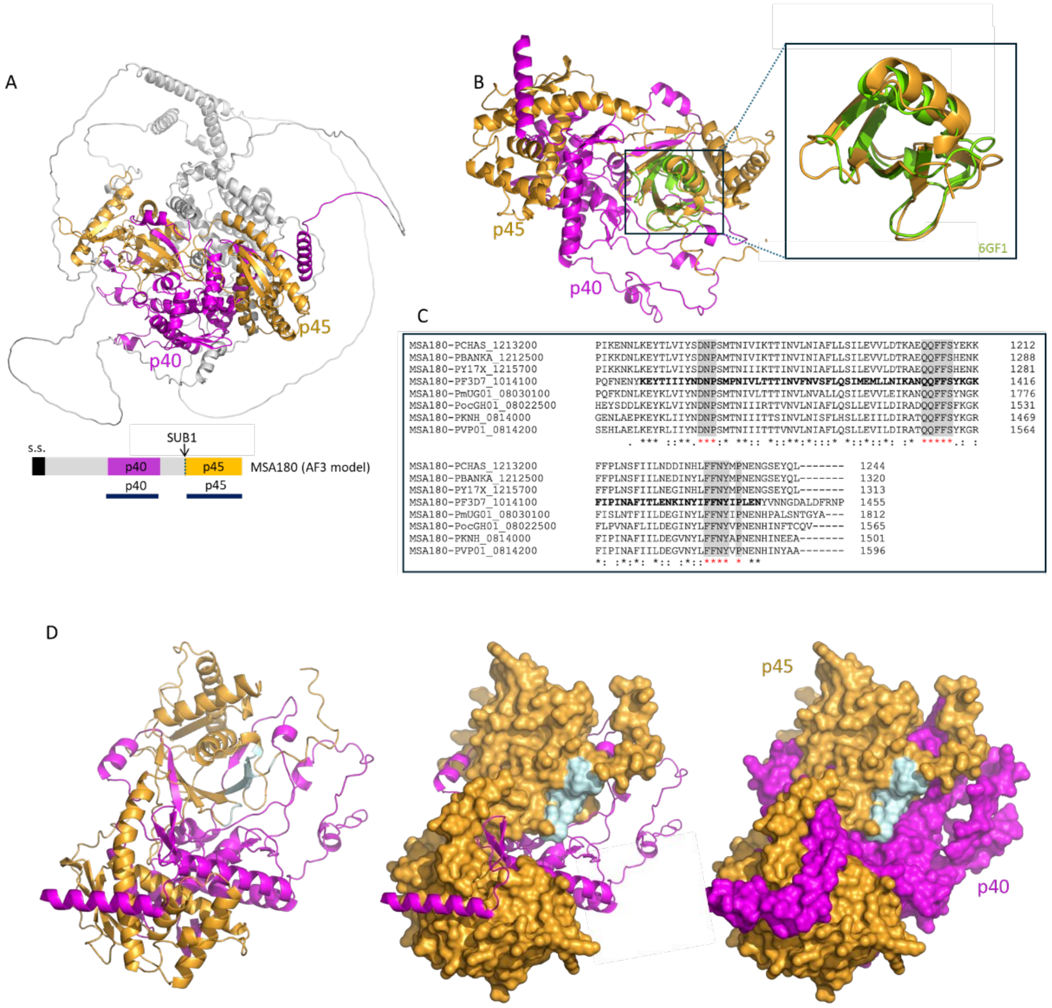


**Figure S1. The SERA6 interacting partner MSA180 contains a conserved putative UBL-domain that is solvent exposed. (**A) Alphafold3 model of MSA180 shown as a grey cartoon with the structured p40 and p45 domains (based on Tan MSY et al. EMBO J 40:e107226, 2021. PMID: 33932049) in magenta and mustard respectively. Note that the p40 and p45 domains interact extensively. Below, schematic of the MSA180 primary sequence with the signal sequence (s.s.) in black and the p40 and p45 segments in magenta and mustard respectively. The known SUB1 processing site is indicated with a black arrow. (B) Isolated MSA180 p40 (magenta) and p45 (mustard) domains. A zoomed view of the C-terminal region of MSA180 p45 is shown, with its fold superimposed onto the ubiquitin-like (UBL) modifier FAT10 (PDB: 6GF1, lemon green cartoon, rmsd 1.9Å) to show their structural similarity. The MSA180 p45 UBL-like domain is not involved in the interaction with MSA180 p40 and is accessible to solvent. (C) Partial primary sequence alignment (ClustalW) spanning the UBL-like region of several

*Plasmodium* MSA180 orthologues to show the degree of conservation in this region. The UBL-like domain of *P. falciparum* MSA180 (PlasmoDB: PF3D7 1014100) is shown in bold. Other sequences are of: *P. chabaudi* (PCHAS_1213200); *P. berghei* (PBANKA_1212500); *P. yoelii* (PY17X_1215700); *P. malari*ae (PmUG01_08030100); *P. ovale* (PocGH01_08022500); *P. knowlesi* (PKNH_0814000); and *P. vivax* (PVP01_0814200). Regions highlighted in grey are completely conserved and are predicted to be surface-exposed on the MSA180 p45 UBL-like domain. (D) Three views of the AlphaFold3 model of the isolated MSA180 p40 and p45 domains, coloured as indicated except that key conserved residues within p45 (shaded grey in alignment C) are shown here in light blue. Most are predicted to be located at the molecular surface of the structure and therefore fully accessible to solvent.

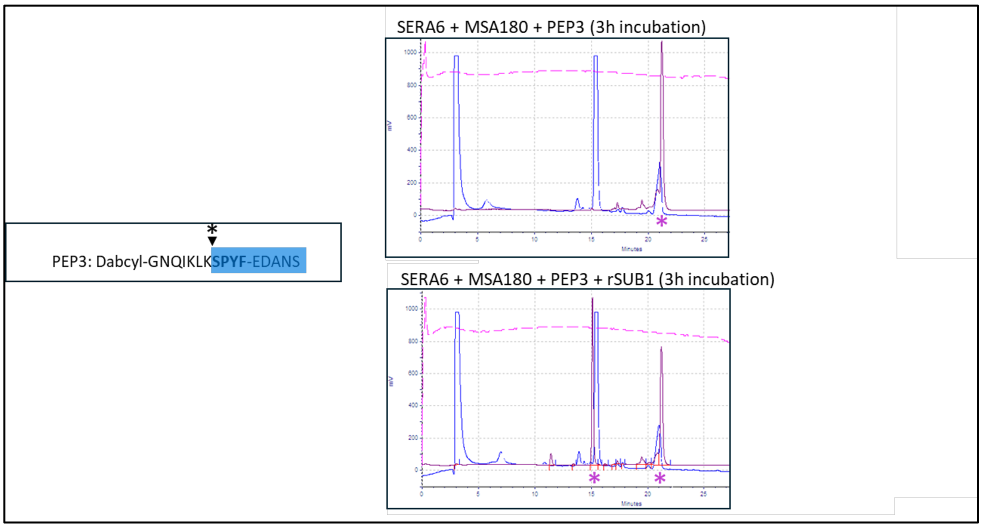


**Figure S2.** ***Ex vivo*** **maturation of SERA6 leads to cleavage of substrate PEP3 at the internal Lys-Ser bond.** Left hand-side: primary structure of fluorogenic peptide substrate PEP3, with the C-terminal region SPYF-EDANS highlighted in blue and the predicted cleavage site asterisked and indicated with an arrowhead. Right-hand side: RP-HPLC elution profiles of samples from the assay described in Fig 4C of the main paper, sampled after a 3 h incubation either in the absence (top) or presence (bottom) of rSUB1. Blue elution profile, UV absorbance (215 nm). Cyan profile, fluorescence (Ex 340 nm, Em 492 nm). The large UV-absorbing peak in both profiles at ~15.5 min is the HA peptide used to elute the HA-tagged FL SERA6 and FL MSA180 proteins during affinity-purification from saponin extracts of schizonts of transgenic *SERA6-mTAP:loxP* and *MSA180-HA3:loxP* parasites respectively. The major fluorescent peak at 21.5 min in the top elution profile (asterisked) corresponds to intact (uncleaved) PEP3, confirmed by electrospray mass spectrometry (expected *mz* 1,793.28, observed 1,792.9). Note that the fluorescence demonstrated by intact PEP3 is probably due to the high levels of acetonitrile in the elution buffer interfering with quenching of the EDANS moiety. The newly appearing fluorescent peak at ~15 min in the lower (+rSUB1) elution profile (also asterisked) was sampled for analysis by electrospray mass spectrometry, detecting a novel product likely corresponding to the PEP3-derived cleavage product SPYF-EDANS (expected *mz* 760.52, observed 760.28).

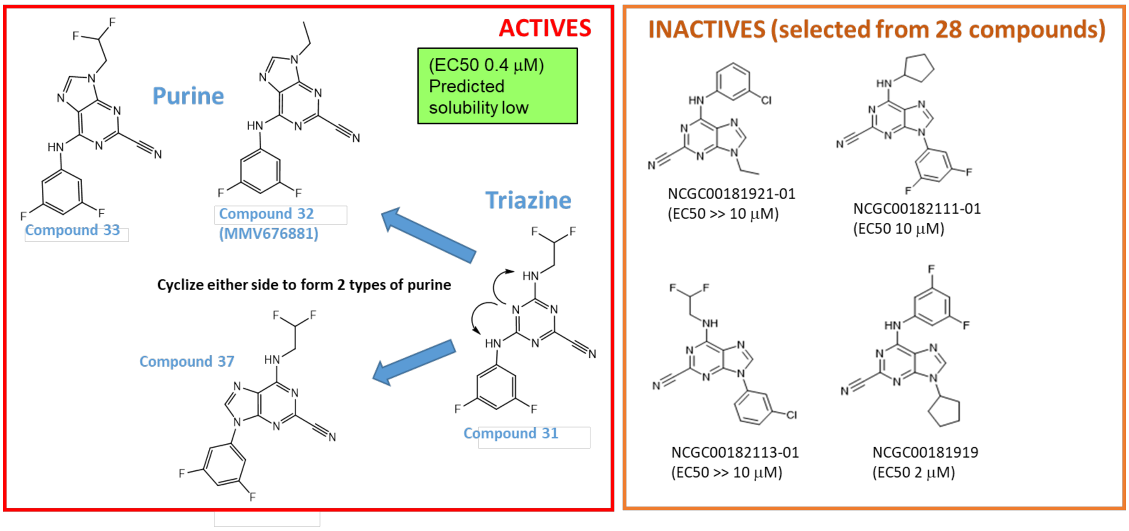


**Figure S3. Structure-activity relationship (SAR) analysis of the NCATS set: the difluorophenyl substituent is important for SERA6 inhibitory potency.** Replacement of the difluorophenyl group in compounds 32 and 37 with a 2-chloro phenyl group abolished potency. Similarly, substitution of the difluoroethyl appendage of compounds 33 or 37 with a cyclopentane ring severely reduced SERA6 inhibitory potency. Also shown (left) is our observation that cyclisation of the triazine compound 31 in two different vectors could rationalise the different architecture of the three purine nitrile actives. Note that the purity and structural integrity of all compounds was confirmed by mass spectrometry. MMV676881 was originally described as compound 32 (see Mott BT et al. J Med Chem 53:52-60, 2010. PMID: 19908842).

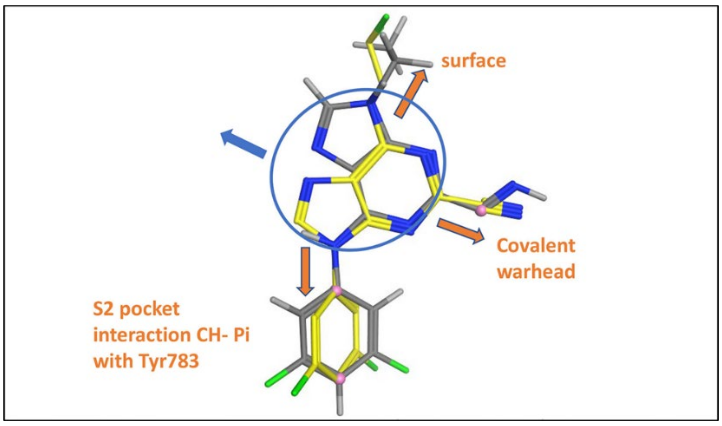


**Figure S4. SAR analysis of the NCATS actives.** Superposition of hit compounds 37 (yellow) and 32 (MMV676881, grey) by selecting the 3 atoms highlighted in pink. Likely key interactions with the Tyr783 in the SERA6 S2 pocket and the solvent-exposed surface are indicated (orange arrows), whilst the purine core packs against the protein (blue arrow). Proposed interactions are based on the x-ray crystal structure of MMV676881 bound to cruzain (PDB: 3I06). Analysis and images generated with MOE 2019.0102.

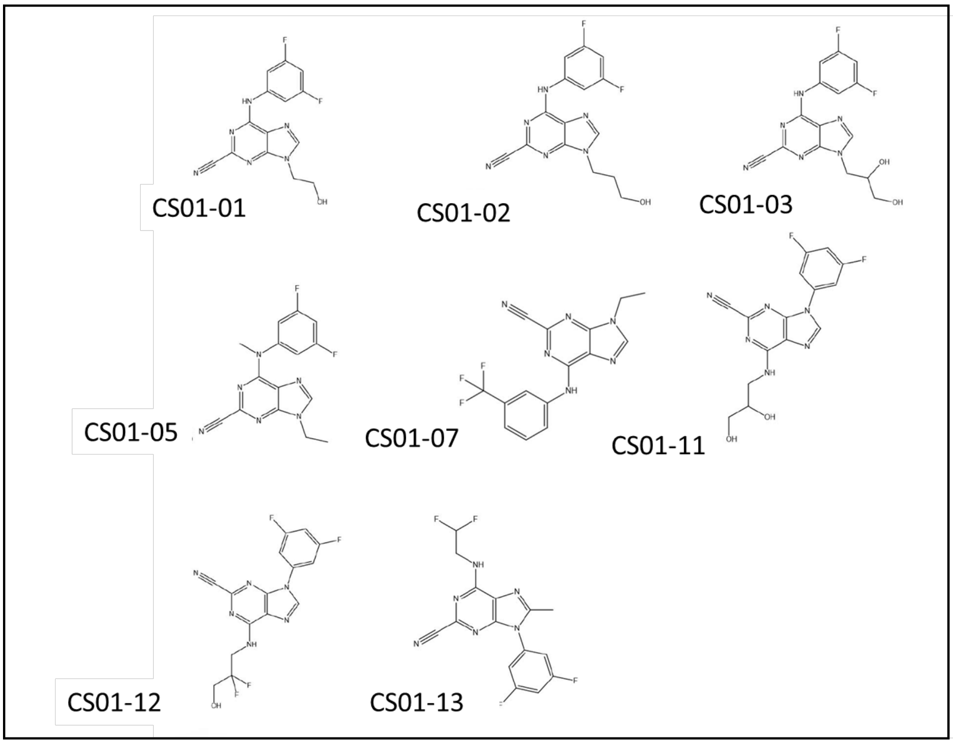


**Figure S5. Molecular structure of the CS compounds.** All compounds were produced by Enamine Ltd and structures confirmed by NMR and LC-MS (liquid chromatography-mass spectrometry.

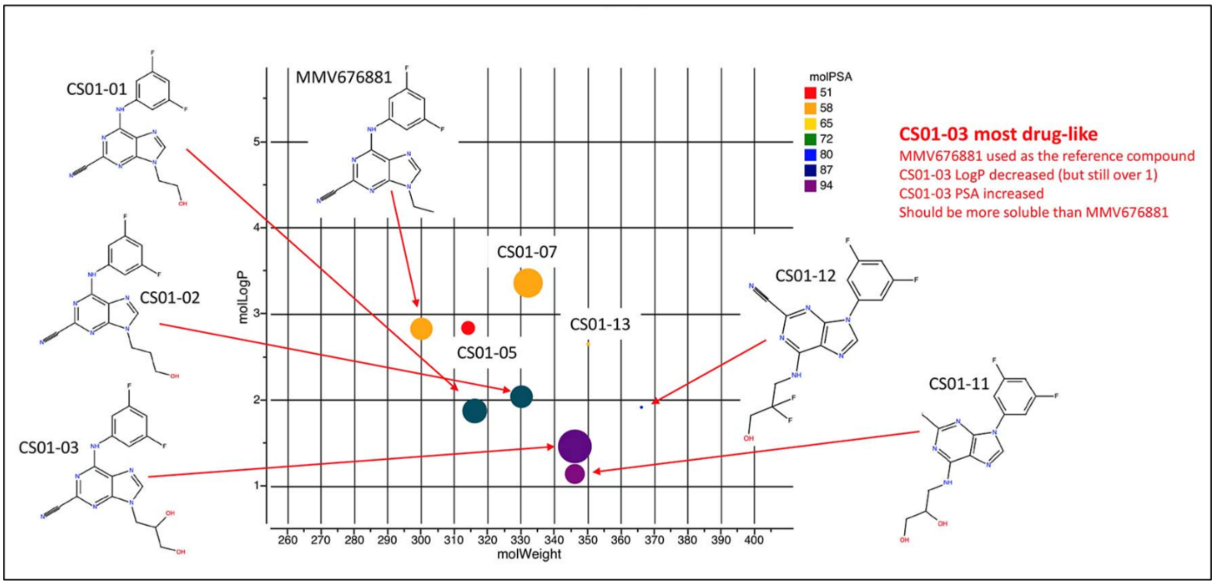


**Figure S6. Predicted chemical properties and drug-likeness of the CS compounds compared to primary hit MMV676881 suggest that CS01-03 is the most drug-like of the CS compound set.** Dot size indicates drug-likeness (the larger the dot, the more drug-like). molLogP, predicted octanol water partition. molPSA, predicted polar surface area (Å$^2$), with dots colour-coded as indicated. Chemical structures of selected compounds are shown. All analysis was performed in ICM-Pro.

| ICM-Pro runs | MMV676881 | CS01-01 | CS01-02 | CS01-03 | CS01-11 | CS01-12 |
|---|---|---|---|---|---|---|
| Run1 (T3) | C1(-17) | C1(-17) | C2(-11) | C1(-22) | C1(-20) | C2(-14) |
| Run2(T3) | C1(-17) | C1(-18) | C1(-17) | C1(-22) | C1(-20) | C2(-13) |
| Run3(T6) | C1(-17) | C1(-18) | C2(-11) | C1(-22) | C1(-19) | C2(-15) |
| Run4(T3) | C1(-17) | C1(-18) | C2(-12) | C1(-22) | C1(-20) | C2(-14) |
| Run5(T4) | C1(-18) | C1(-18) | C1(-16) | C1(-22) | C1(-20) | C1(-17) |

C1: conformation 1
C2: conformation 2
T: thoroughness or length of simulation, varied from 3 (standard) to 6
ICM docking scores in brackets, the lower the score, the better the docking compound

**Figure S7. Covalent docking analysis of the purine nitrile hits shows that CS01-03 (yellow star) displays the best docking score.** Results are shown from 5 independent docking analyses in ICM-Pro using the SERA6 model. Docking scores are shown in parentheses. C1 and C2 refer to the 2 different docking conformations observed in the case of CS01-02 and CS01-12.

| STRUCTURE | Isomer | ICM-Pro run1 T6C1 scores | ICM-Pro run2 T10C1 scores | ICM-Pro run3 T4C1 scores | ICM-Pro run4 T5C1 scores | ICM-Pro run5 T7C1 scores |
|---|---|---|---|---|---|---|
| $C_{15}H_{12}F_2N_6O_2$ 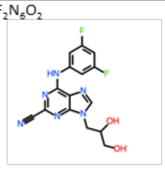 | CS01-03 R ISOMER | -22 | -22 | -22 | -22 | -22 |
| $C_{15}H_{12}F_2N_6O_2$ 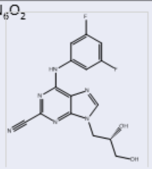 | CS01-03 S ISOMER | -9 UD | -12 UD | -9 UD | -12 UD | -12 UD |

T: thoroughness, length of simulation
C: number of conformations
UD: ‘upside down’ conformation

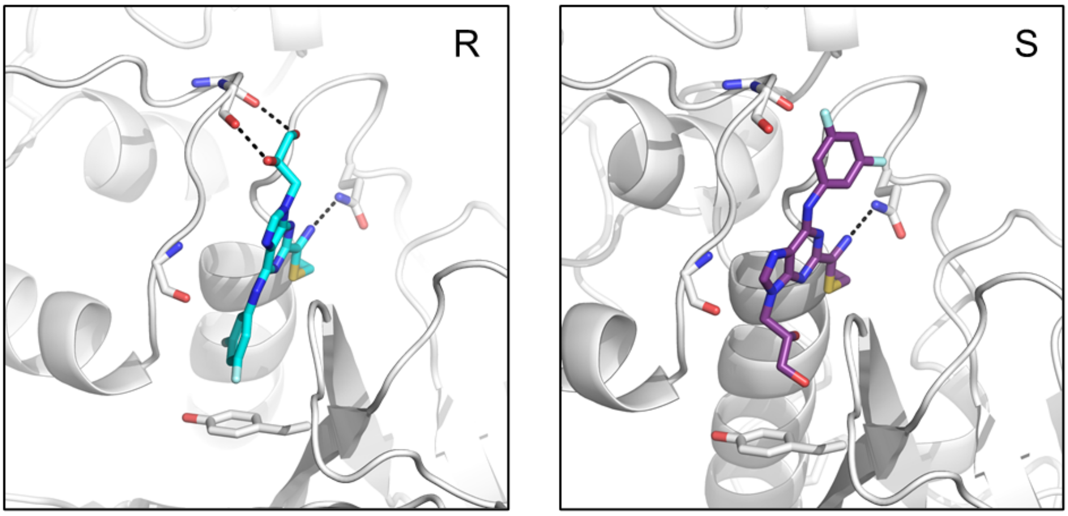


**Figure S8. Docking analysis suggests a preference for binding of the CS01-03 R-enantiomer to SERA6.** Top, table showing the results of ICM-Pro covalent docking analysis of the two enantiomers of CS01-03 into the AlphaFold SERA6 model. Whereas the S-enantiomer gave poor docking scores and only then in an ‘upside-down’ (UD) conformation in which the difluorophenyl group faces solvent, the R-enantiomer consistently gave much better docking scores predictive of high affinity binding. Bottom, structures of the enantiomers docked into the SERA6 S2 pocket.

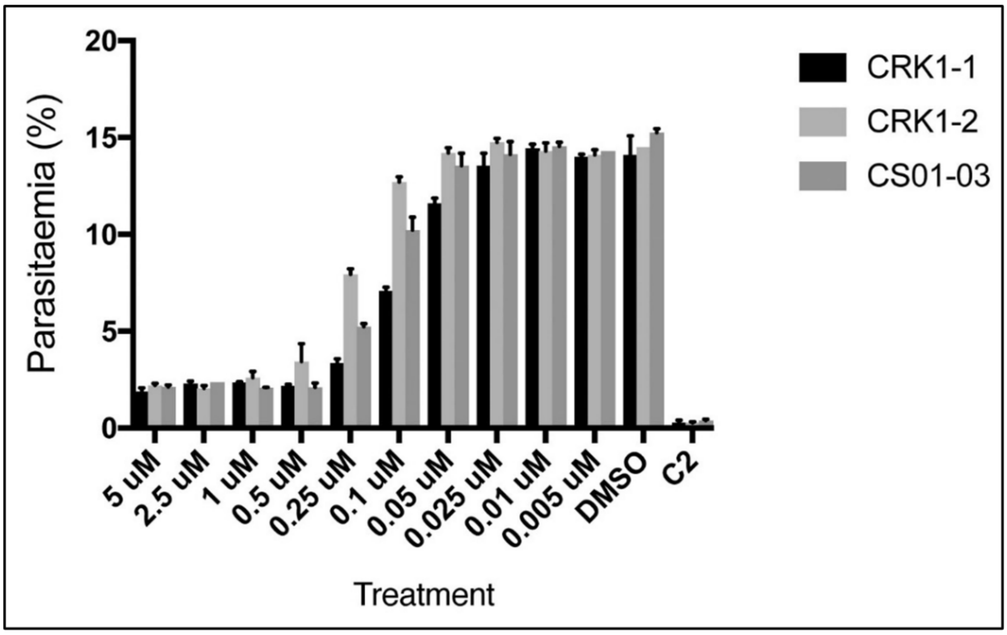


**Figure S9. Anti-parasite activity of the separated CS01-03 enantiomers.** Dose-response in the *P. falciparum* schizont-to-ring transition assay for compound CS01-03 and its separated enantiomers CRK1-1 and CRK1-2 (note that it was not established which of these corresponded to the S- and R-enantiomers). The estimated $EC_{50}$ for CRK1-1 was ~100 nM whilst CRK1-2 was 3-4-fold less potent. Unfractionated CS01-03 displayed intermediated potency as expected. Vehicle only (DMSO, 1% v/v) and the PKG inhibitor C2 (1 μM) were included as negative and positive controls respectively. Results shown represent mean values from triplicate assays (error bars, SD).

| Cpd | Solubility (μM) | CHI LogD | MCLint | |
|---|---|---|---|---|
| | | | mL/min/mg protein | MCLint (mL/min/g liver*) |
| CRK1-1 | 10.0 | 2.25 | 0.133 | 6.97 |
| CRK1-2 | 22.6 | 2.25 | 0.078 | 4.1 |

*Using a scaling factor of 52.5 mg microsomal protein per g liver

**Figure S10. In vitro drug metabolism and pharmacokinetic (DMPK) analysis of CS01-03 enantiomers CRK1_1 and CRK1_2 indicates good solubility and stability for both enantiomers.** Solubility values greater than 10 μM are important for preclinical testing. CHI LogD, an experimental measure of lipophilicity. MCLint, intrinsic clearance in human microsomes. Solubility, CHI LogD and microsomal stability assays were kindly performed by Jennifer Riley, Drug Discovery Unit, University of Dundee, UK.

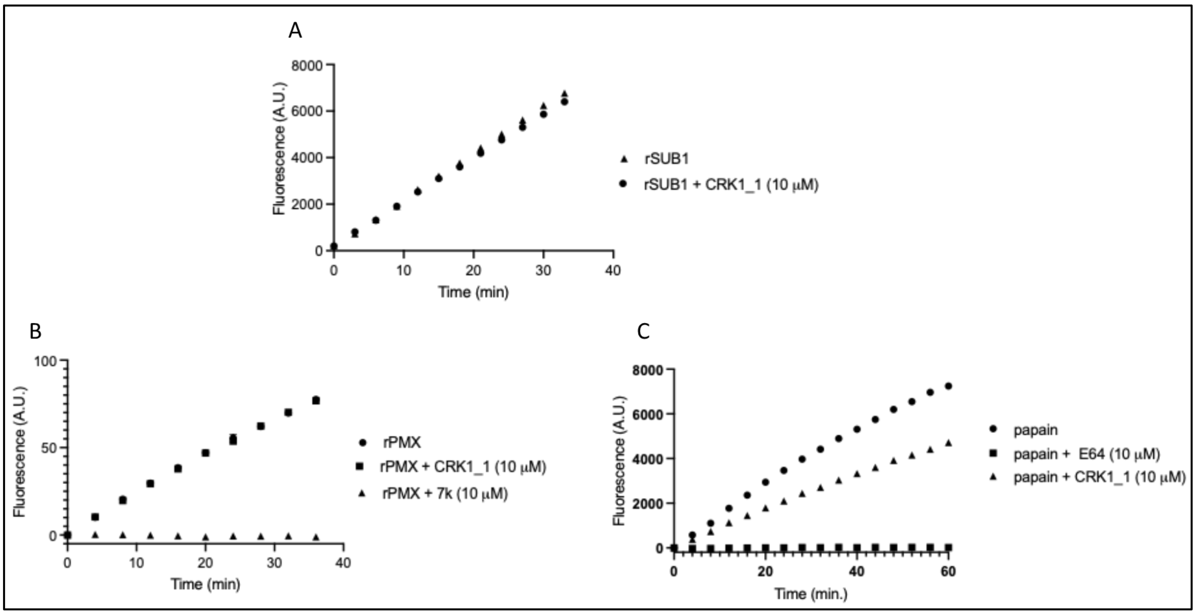


**Figure S11. Selectivity of nitrile compound CRK1_1.** The compound was tested against: (A) rSUB1; (B) recombinant PMX; and (C) recombinant papain, as described in the main manuscript. Progress curves show fluorescence increase over time as the relevant fluorogenic substrates are cleaved. CRK1_1 showed no inhibition of either rSUB1 or PMX at the concentration used (10 μM), and only partial inhibition of papain. Positive control inhibitors used in the PMX and papain assays were compound 7k (Kovada V et al J Med Chem 66:10658-10680, 2023. PMID: 37505188) and the epoxide E64 respectively.

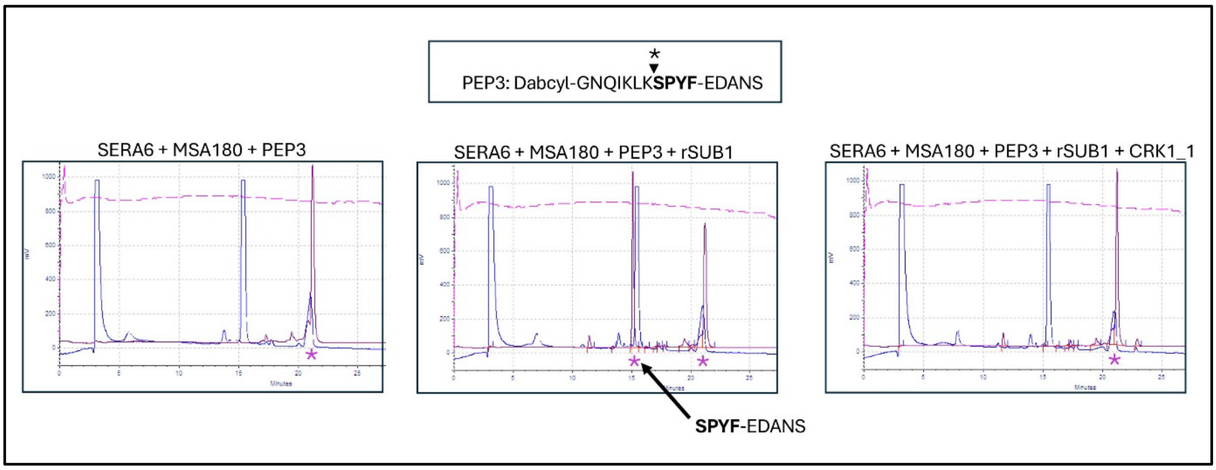


**Fig S12. Compound CRK1_1 directly inhibits SERA6 peptidase activity.** RP-HPLC analysis of supernatants from the *in vitro* PEP3 cleavage assay shown in Fig 7C of the main manuscript shows that formation of the fluorescent SPYF-EDANS cleavage product (indicated) is blocked by the presence of CRK1_1.
